# Cancer phylogenetic tree inference at scale from 1000s of single cell genomes

**DOI:** 10.1101/2020.05.06.058180

**Authors:** Sohrab Salehi, Fatemeh Dorri, Kevin Chern, Farhia Kabeer, Nicole Rusk, Tyler Funnell, Marc J Williams, Daniel Lai, Mirela Andronescu, Kieran R. Campbell, Andrew McPherson, Samuel Aparicio, Andrew Roth, Sohrab Shah, Alexandre Bouchard-Côté

**Author notes:** Equal contribution (Sohrab Salehi, Fatemeh Dorri).

## Abstract

A new generation of scalable single cell whole genome sequencing (scWGS) methods allows unprecedented high resolution measurement of the evolutionary dynamics of cancer cell populations. Phylogenetic reconstruction is central to identifying sub-populations and distinguishing the mutational processes that gave rise to them. Existing phylogenetic tree building models do not scale to the tens of thousands of high resolution genomes achievable with current scWGS methods. We constructed a phylogenetic model and associated Bayesian inference procedure, sitka, specifically for scWGS data. The method is based on a novel phylogenetic encoding of copy number (CN) data, the sitka transformation, that simplifies the site dependencies induced by rearrangements while still forming a sound foundation to phylogenetic inference. The sitka transformation allows us to design novel scalable Markov chain Monte Carlo (MCMC) algorithms. Moreover, we introduce a novel point mutation calling method that incorporates the CN data and the underlying phylogenetic tree to overcome the low per-cell coverage of scWGS. We demonstrate our method on three single cell datasets, including a novel PDX series, and analyse the topological properties of the inferred trees. Sitka is freely available at https://github.com/UBC-Stat-ML/sitkatree.git.

## 1 Introduction

A main challenge in investigating cancer evolution is the need to resolve the subpopulation structure of a heterogeneous tumour sample. Advances in next generation scWGS have enabled more accurate, quantitative measurements of tumours as they evolve [1, 2, 3, 4]. Phylogenetic reconstruction is central to identifying clones in longitudinal xenoengraftment [5, 6] as well as patients [7], and has been used to approximate the rate and timing of mutation [8] to determine the origins and clonality of metastasis [9, 10].

Single cell cancer phylogenetics is an evolving field. Multiple approaches, spanning different study designs and data sources are reviewed in [11]. Many phylogenetic inference methods such as Scite, SciΦ, OncoNEM and SciClone use the often-made infinite site model assumption and consider point mutations as input or assume a small number of leaf nodes [12, 13, 14, 15]. However, emerging single cell platforms produce up to thousands of single cell genomes and are suitable for determining copy number aberrations (CNA) [16, 1]. Compared to phylogenetic methods based on point mutations, fewer can build phylogenies from large scale CN data. Recent related work includes MEDALT [17], which models single-cell copy number lineages using a spanning tree over cells rather than a phylogeny; and SCARLET [18], which proposes a point mutation-based phylogeny inference procedure that calibrates mutation losses with copy number profiles. Distance based and agglomerative clustering methods such as neighbour joining [19, 20] are often used to elucidate hierarchical structures over cells, in particular, in the context of CNA, see [17], however, distance-based methods tend to produce less accurate tree reconstructions [21, 22, 23, 24].

Single cell whole genome DNA sequencing provides low, yet uniform coverage [1]. That is, the sequenced reads cover for each single cell about 0.1 per cent of the genome. In this setting, most point mutations are not observed in most single cells making calling SNVs difficult. However, it is possible to identify the relative copy number for segments of the genome. These copy number events can be used as phylogenetic markers.

We describe sitka, a phylogenetic model and an associated Bayesian inference procedure designed specifically for inference based on CN information extracted from scWGS data (see **Fig. 3**). Our method addresses two key challenges: first, each CNA event typically affects a large number of genomic sites, breaking the independence assumptions required by existing phylogenetic methods [25, 15, 13, 26]; second, while detailed modelling of dependent evolutionary processes is in principle possible, they entail computational requirements incompatible with the scale of modern scWGS data [27]. To confront these two difficulties, sitka uses a novel phylogenetic encoding of CN data, providing a statistical-computational trade-off by simplifying the site dependencies induced by rearrangements, while still forming a sound foundation to phylogenetic inference. Based on this encoding, we propose an innovative phylogenetic tree exploration move which makes the cost of Markov chain Monte Carlo (MCMC) iterations bounded by *O*(|*C*| + |*L*|), where |*C*| is the number of cells and |*L*| is the number of loci. In contrast, existing off-the-shelf likelihood-based methods incur an iteration cost of *O*(|*C*| |*L*|) [28, 13, 15]. Moreover, the novel move considers an exponential number of neighbouring trees whereas off-the-shelf moves consider a polynomial size set of neighbours.

Sitka’s workflow proceeds by partitioning the CN information extracted from scWGS data of each cell into bins of fixed size (500Kb) with an integer CN state associated with each bin. This input data is then transformed into a binary format that captures CN changes, but not their direction or magnitude. Conditional on this binary encoding, sitka then yields an approximate posterior distribution on compatible phylogenetic trees.

Potential applications of sitka include lineage tracing and subclonal structure identification. In lineage tracing the goal is to relate single cells to their ancestors based on genomic markers. This is especially useful in experimental designs where multiple samples from the same subject exist, e.g., multi-region or timeseries studies [5]. In subclonal structure identification, the topology of the inferred phylogeny can be used to make inferences about the evolutionary forces acting on the trees [29].

We compare sitka with other tree inference methods on three real-world datasets, including triple negative breast cancer patient derived xenograft samples, high grade serous ovarian primary and matched relapse samples. Since the true phylogeny is unknown, we design a phylogenetic goodness-of-fit framework to quantitatively assess the performance of our method and to visualize reconstruction confidence as well as violations of our assumptions.

We use the sitka inferred trees to analyse the topological properties of the real-world datasets. Finally, we introduce a model extension that enables the placement of single nucleotide variants (SNV) with high levels of missingness on a tree inferred from the CN data.

## 2 Results

### 2.1 Sitka: scalable single cell phylogenetic tree inference

**Fig. 3** shows the workflow of the sitka method. Sitka is based on a transformation of single cell copy number matrices retaining only presence or absence of changes in copy number profiles between contiguous genomic bins. This transformation allows us to approximate a complex evolutionary process (integer-valued copy numbers, prone to a high degree of homoplasy and dense dependence structure across sites) using a probabilistic version of a perfect phylogeny (see **Fig. 1**). We leverage the special structure created by the change point transformation to build a special purpose MCMC kernel, which has better computational scalability per move compared to classical phylogenetic kernels (Methods section 9.4.3).

**Figure 1.**
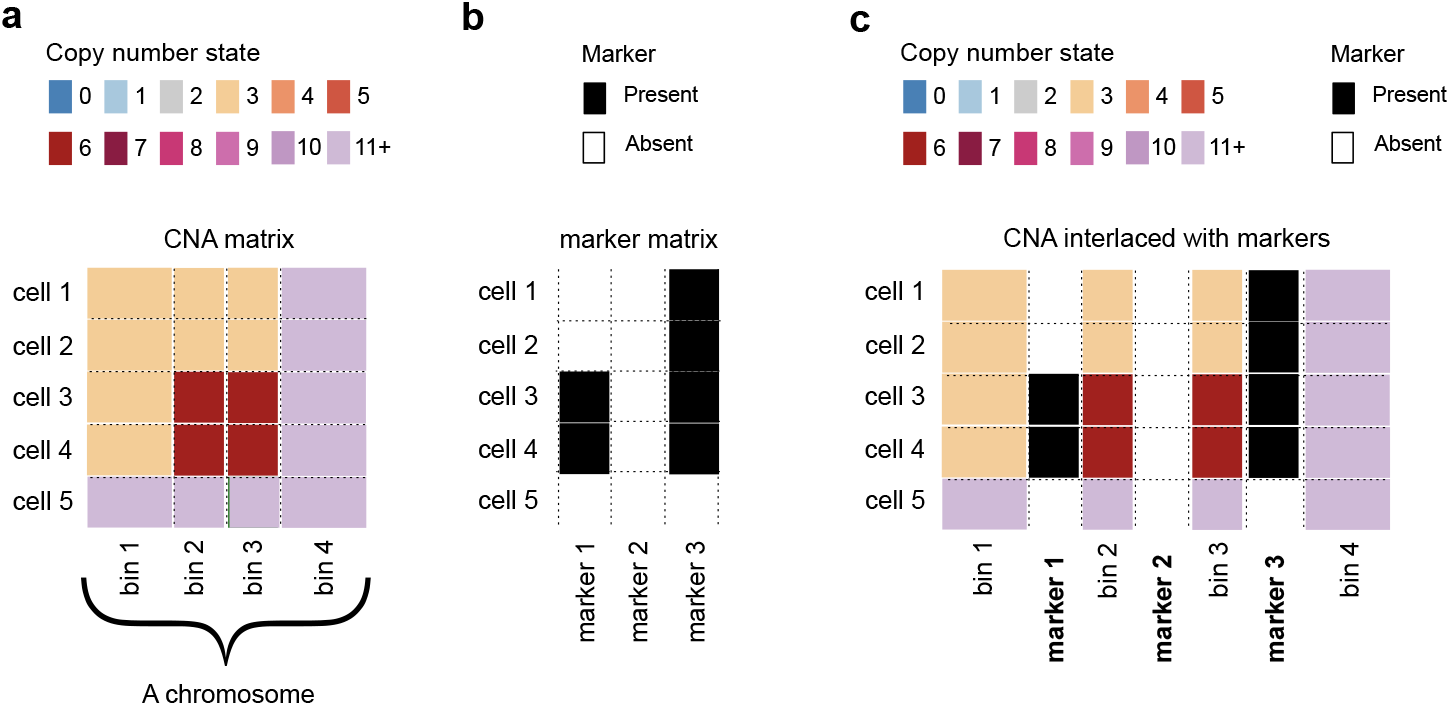
Description of the process involved in the construction of *markers*, the input to the sitka model. A *bin* is a contiguous set of genomic positions. Each pair of consecutive bins (e.g., bins 1 and 2 in (**a**)) is associated with a *marker* (e.g., marker 1) that measures for each individual cell, whether there is a difference between the CNA states of the two bins. (**a**) The observed CNA matrix for a subset of bins on a chromosome. The rows are sequenced single cells, and the columns are bins. The CN states are colourcoded. (**b**) The three markers shown are associated with the four bins. Each marker records the presence (black) or absence (white) of a CN state change between a pair of consecutive bins. Note that in the CNA matrix, there is a CN change at row 3 from bin 1 to bin 2 (CN state 3 to 6). This is reflected in the marker matrix, at row 3 of marker 1 with a black square. There are no changes between bins 2 and 3 across any rows in the CNA matrix. This is reflected in marker 2 comprising all white squares. (**c**) For visualisation purposes, the CNA matrix can be interlaced with the marker matrix to more clearly show where the CNA changes occur. Each column of the marker matrix is inserted between the associated pair of columns in the CNA matrix. The resulting matrix is an example of an *augmented* view that combines data from two or more sources (here the CNA matrix and the marker matrix). In an augmented view, we call columns from each source a *channel*.

**Figure 2.**
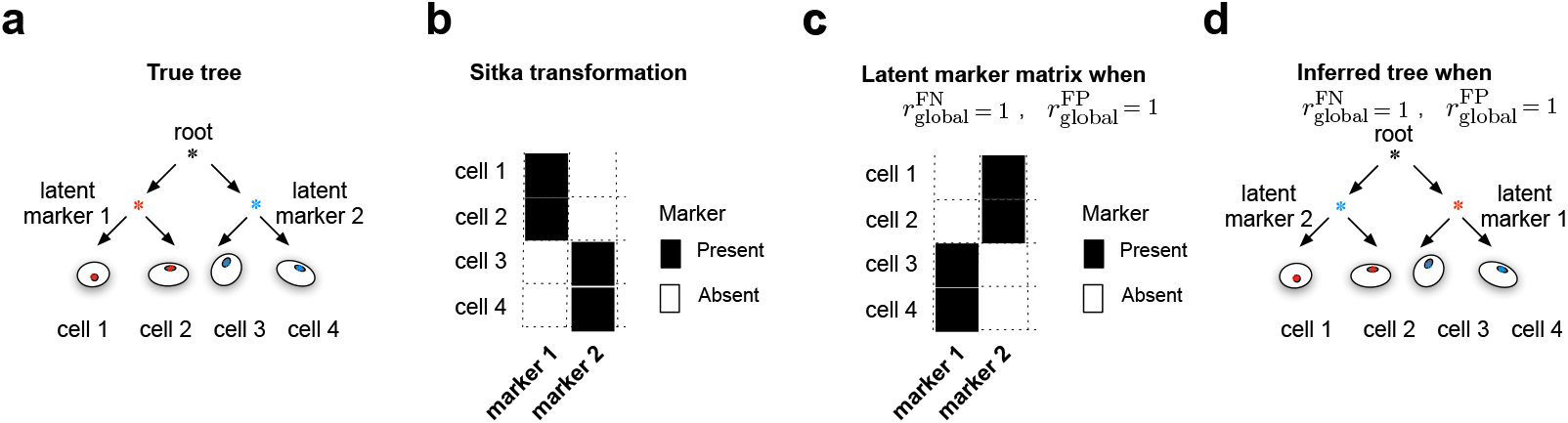
Pathological tree reconstruction under default observation prior. (**a**) The true tree reconstruction in a simple example with a balanced phylogeny with two clades of size two, and two unique markers, coloured red and blue, that distinguish the left and right clades respectively. (**b**) The binarised input matrix corresponding to the four cells at the two markers. The desired observation error rates should be zero and the latent and observed marker matrices should match exactly, as the perfect phylogeny assumption holds. If the observation error parameters are set to one, that is 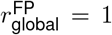 and 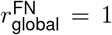, then the latent marker matrix with all entries flipped as shown in (**c**) will have an equal likelihood under this setting as the desired latent matrix has when error rates are set to zero. (**d**) The incorrect tree reconstruction where the left and right clades are erroneously assigned to the blue and red markers.

**Figure 3.**
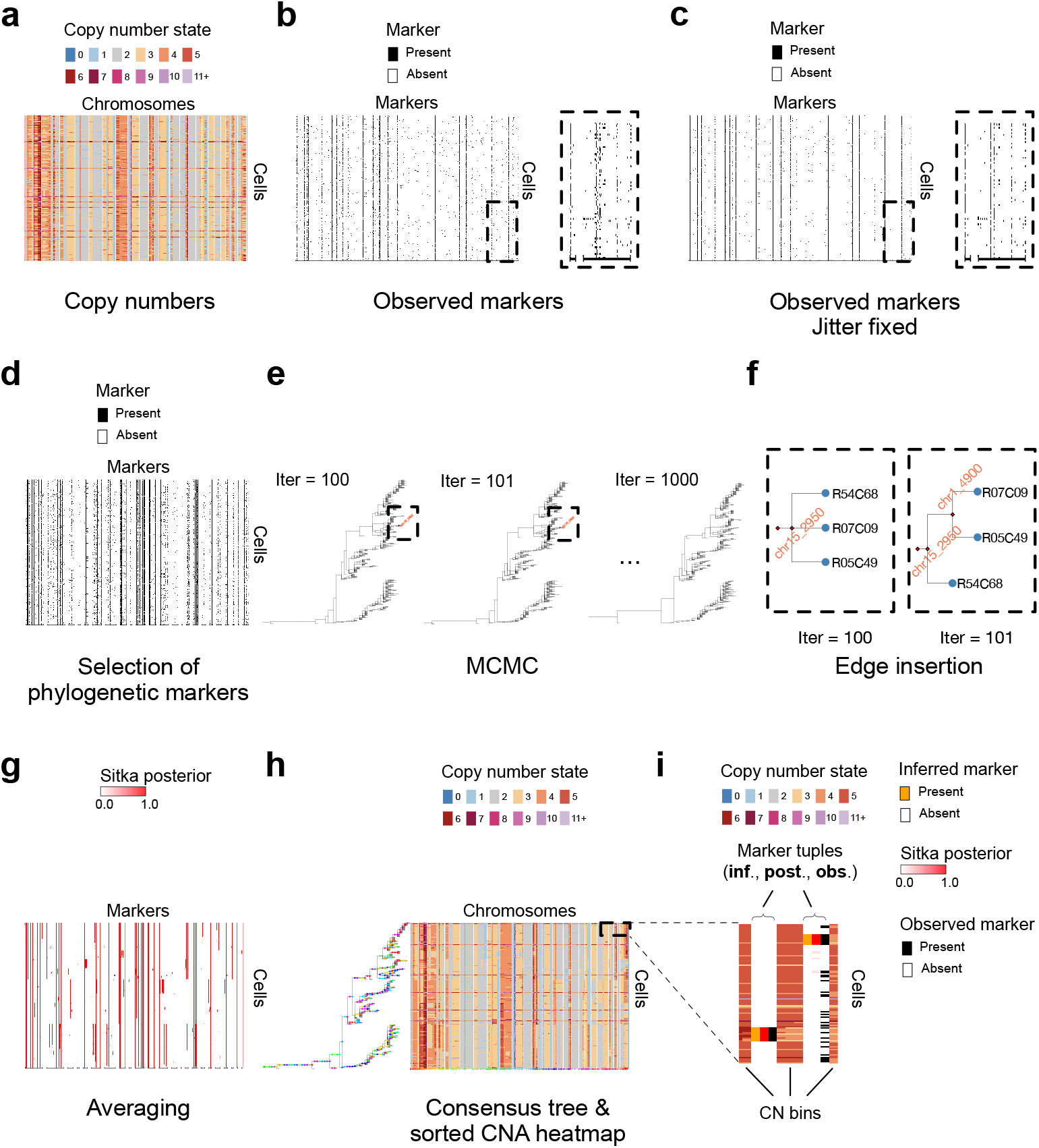
Workflow of sitka. (**a**) Sitka takes copy number calls data from a heterogeneous single-cell population. The cells (rows of the copy number matrix) are randomly sorted. (**b**) A lossy binary transformation is applied to obtain markers data. (Methods section 9.2 and **Fig. 1**). Note that each single-cell is now represented by the presence or absence of CN changes between consecutive bins. (**c**) The boundary conditions are smoothed to account for cell-specific marker misalignment. (Methods section 9.3) to correct for this marker misalignment. Note how the columns in the inset in panel-**c** are less noisy than their counterpart in panel-**b**. (**d**) A subset of markers present in at least 5 percent of the cells are chosen for input to the tree inference algorithm. (**e**) An MCMC algorithm efficiently explores the tree space. (**f**) An example of an edge-insertion. The two insets are zoomed in from panel-**e**. Each inset depicts a subtree, where red diamonds and blue circles denote marker nodes and single-cells respectively. Also see **Supplementary Fig. 1** for details. (**g**) The indicator matrix of all post-burn-in MCMC trees are averaged to generate a matrix indicating the posterior probability of a cell being attached to a marker (Methods section 9.4.5). (**h**) The copy number data in (**a**) is sorted according to the inferred consensus tree, shown on the left of the matrix. (**i**) The inset shows the tuple of marker columns in the context of the copy number calls, namely **inf**. (inferred markers, i.e., latent state *x*_*c,l*_), **post**. (posterior probability of the latent state *x*_*c,l*_), and **obs**. (observed markers), interlaced with the CN columns (similar to **Supplementary Fig. 1**). The results are from the *SA*535 dataset, a triple negative breast cancer patient derived xenograft sample (Methods section 2.2).)

We visualise the input data to sitka in a colour-coded matrix exemplified in **Supplementary Fig. 1-a**. Each row in the matrix corresponds to an individual cell that has been sequenced in a single-cell platform. Each column in the matrix is a locus that is represented by a bin (a contiguous set of genomic positions). We assume that the integer copy number of each bin has been estimated as a preprocessing step, e.g., using a hidden Markov model [16]. In **Supplementary Fig. 1-a** the copy number state is encoded by the colour of each entry in the matrix.

The output of sitka includes two types of directed rooted trees. Type I is the tree used for MCMC sampling in the inference procedure, and type II, which is derived from type I, is used in visualisation (**Fig. 4-a-c**). The set of nodes in a type I tree is given by the union of the cells, the CN change points (markers) under study, and a root node *v*\*. The topology of a type I tree bears the following phylogenetic interpretation: given a cell *c* in the tree, *c* is hypothesized to harbour the markers in the shortest path between *c* and the root node *v*\*, and only those markers. We enforce the constraint that all cells are leaf nodes, while markers can be either internal or leaf nodes. Markers placed at the leaves are interpreted as outliers, for example measured CN change points that are false positives.

**Figure 4.**
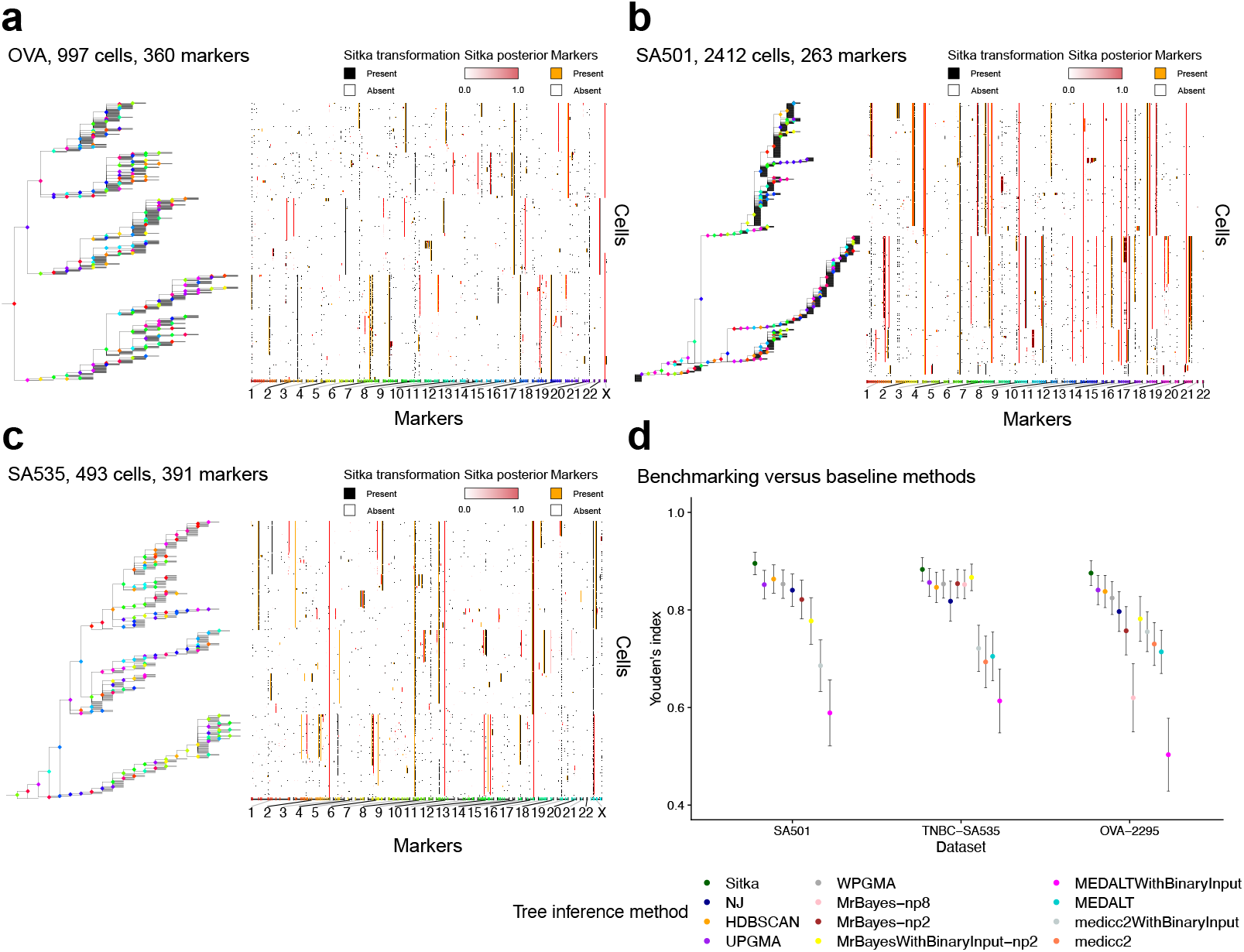
Results over real-datasets and benchmarking against baseline methods. (**a**), (**b**), and (**c**) show the consensus tree and marker-space matrix for the *OV A, SA*501, and *SA*535 datasets respectively. (**d**) Comparison to baseline methods.)

To convert a type I tree to a type II tree, we remove from the type I tree all marker nodes that are leaf nodes, i.e., markers that are not present in any cells. We also collapse into a single node, the list of connected marker nodes that have exactly one descendent (i.e., chains). **Fig. 5** shows a small *type I tree*, its transformation to a *type II tree* and the respective marker matrix. We visualise the input matrix and the estimated tree simultaneously by sorting the individual cells (rows of the matrix) such that they line up with the position of the corresponding leaves of the tree.

**Figure 5.**
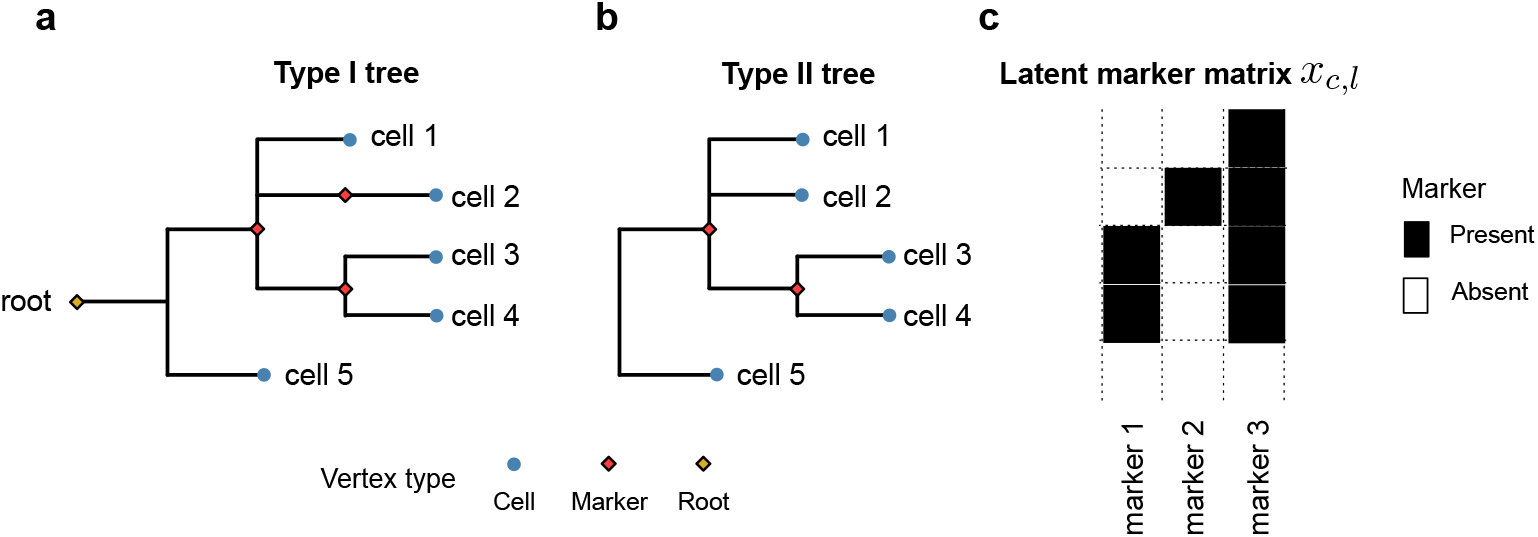
Visualisation of a small tree and its corresponding marker matrix. A small type I tree *t* (**a**), its transformation into a type II tree (**b**), and the corresponding marker matrix *x* = (*x*_*c,l*_) (**c**). The red nodes in (**a**) correspond from top to bottom to markers 2, 3, 1 in this order. Given a tree *t*, the latent marker matrix *x* is a deterministic function *x* = *x*(*t*). Note that the clade comprising single-cells 3 and 4 has support in both markers 1 and 3. For clarity, we do not visualise type I trees, but plot their transformation, i.e., type II trees as follows. We remove from the type I tree all marker nodes that have *x*_*c,l*_ = 0 for all single-cells *c*. Lists of connected edges that have exactly one descendent (i.e., chains) are also collapsed into a single edge, e.g., the edge corresponding to markers 2 and 3 are collapsed into one edge (since marker 2 has only one descendent, namely single-cell 2).

Sitka uses change points as phylogenetic traits modelled using a relaxation of the perfect phylogeny assumption. For a phylogenetic tree, the perfect phylogeny assumption holds if and only if for all markers *l, l* changes at most once from its ancestral state over the tree. Change points arising from non-overlapping CNA events (i.e., such that the genomic locations affected by the CN event do not intersect) preserve the perfect phylogeny assumption. **Fig. 6** shows examples of overlapping CNA events and their effect on markers. The two scenarios that can lead to the violation of the perfect phylogeny assumption are (i) when a CNA gain event is followed by an overlapping loss event or (ii) when a loss event is followed by an overlapping loss event, and the second event removes either end-point of the first event. For both (i) and (ii), a violation occurs only when the second overlapping event hits the same copy as the first event.

**Figure 6.**
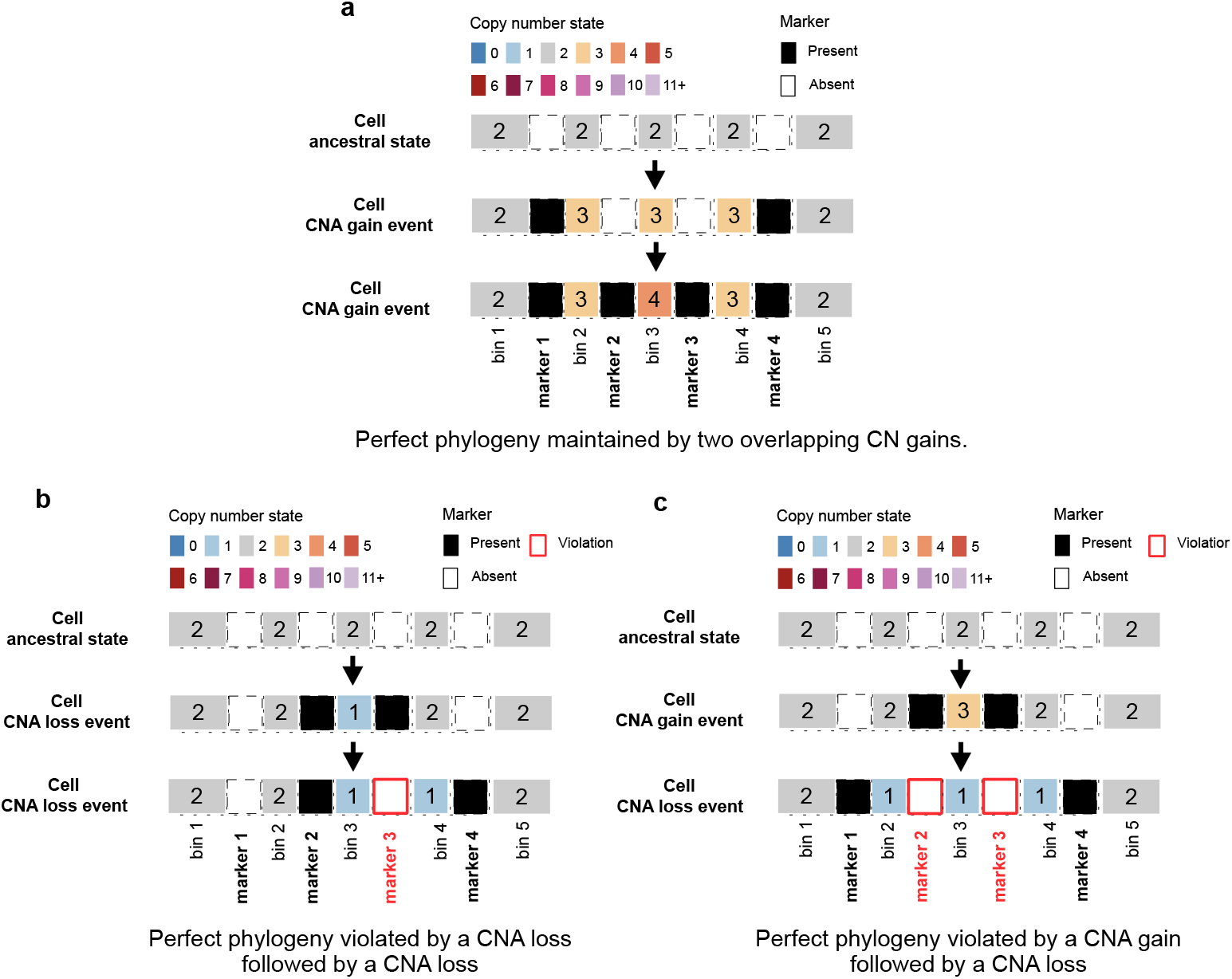
Perfect phylogeny and effects of overlapping CNA events. The effects of overlapping CNA events on the perfect phylogeny assumption. A segment of a chromosome with five consecutive bins and their four corresponding markers are shown. Each panel follows the CN states interlaced with markers for a cell at the ancestral state (top), after a CNA event (middle), and after a second overlapping CNA event (bottom). The numbers in the CNA squares show the integer CN state (e.g., the ancestral state has two copies of the 5-bins long segment). (**a**) Two overlapping CNA gains maintain the perfect phylogeny assumption. If the infinite site assumption holds, it is unlikely for the end-points of the two gain events to exactly match. The same argument holds for a CNA loss followed by a CNA gain event. Note that in these cases, once a change point is acquired, it is not lost. (**b**) If a loss event is followed by another loss event in which either end-points of the first event is removed, the perfect phylogeny assumption will be violated (e.g., marker 3 is lost after the second loss event). Note that a violation does not occur if the loss events hit different copies of a segment. (**c**) Similarly, if a gain event is followed by a loss event, only if the latter erases the end-points of the former is the perfect phylogeny violated. Note how marker 2 and marker 3 are lost after the second CNA event.

Imposing a perfect phylogeny on the *observed* change points is restrictive, as we expect both violations of the assumptions (e.g., due to homoplasy), and measurement noise. To address this we use an observation model (Methods section 9.4.1) which assigns positive probability to arbitrary deviations from the perfect phylogeny assumption, while encouraging configurations where few markers and cells are involved in violations. Subsequently we impose the perfect phylogeny assumption on a *latent* maker matrix defined as follows. Given a type I tree *t*, the latent marker matrix *x* is a deterministic function *x* = *x*(*t*). We compute *x* : *t* → {0, 1} ^*C*×*L*^ by setting *x*_*c,l*_ = 1 if the single-cell *c* is a descendent of the marker node *l* in tree *t*, and otherwise *x*_*c,l*_ = 0. We use *y*_*c,l*_ to refer to the observed change point *l* in individual cell *c* (Methods section 9.4.1).

Synthetic experiments show that sitka’s performance decreases roughly linearly as a function of the rate of the key types of expected violation of the perfect phylogeny assumption (**Fig. 7-a,b**, Methods section 9.5).

**Figure 7.**
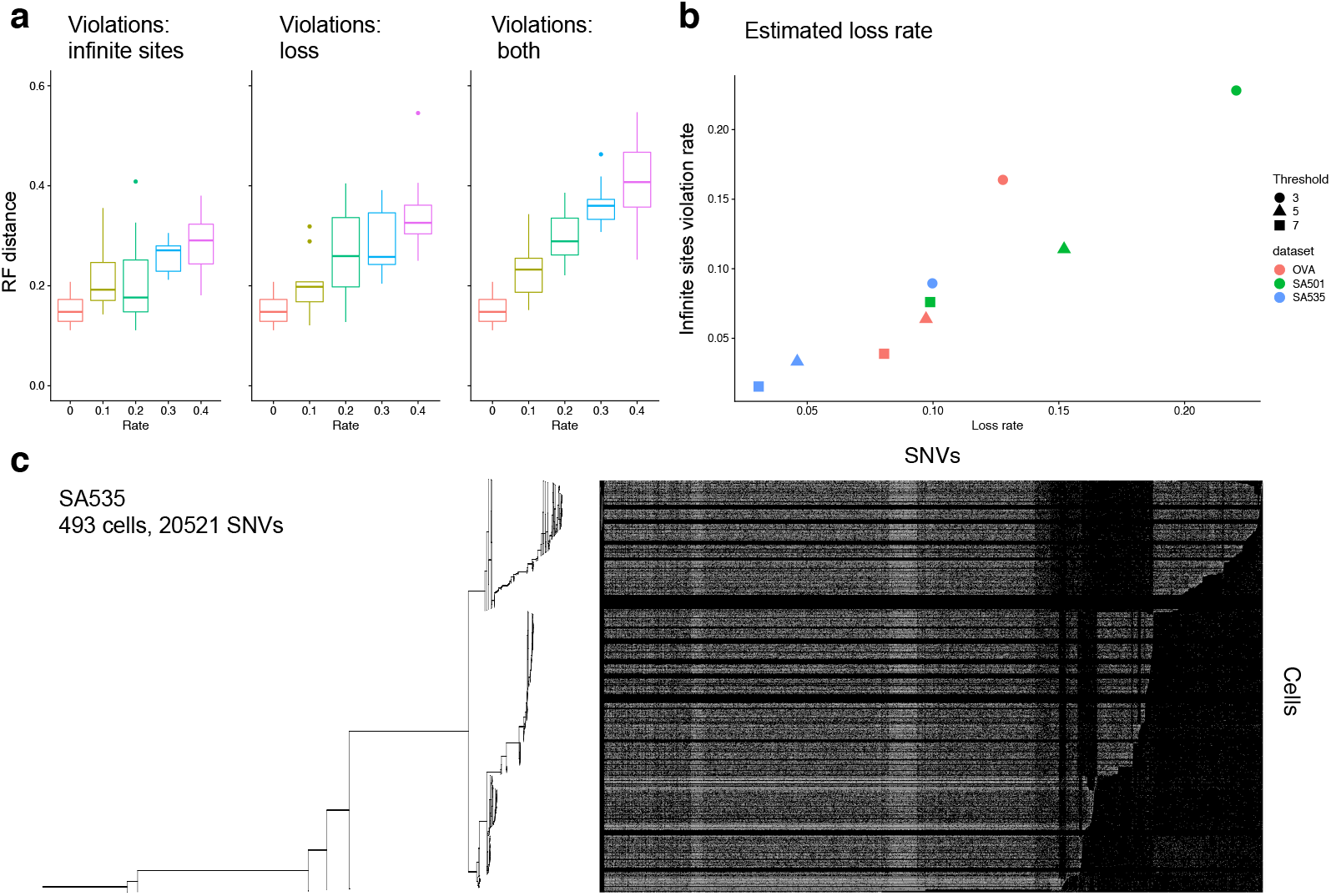
Synthetic experiments and an application to point mutation placement. (**a**) RF distance of sitka tree estimate to the best-possible tree. The first plot holds *p*_*is*_ constant at zero. The second plot holds *p*_*loss*_ constant at 0. The third plot varies *p*_*is*_ = *p*_*loss*_ jointly. **(b)** Estimation of violation rates in real data and a set of synthetic data. (**c)** Over 20,000 SNV’s with high levels of missingness are placed on a backbone tree inferred from the CNA data for *SA*535.

### 2.2 Performance of sitka relative to alternative approaches

We compare the performance of sitka to alternative approaches (Supplementary section 2) on three scWGS datasets introduced here (**Fig. 4-a-c**). The first dataset, *SA*535, is generated for this project and contains 679 cells from three passages of a triple negative breast cancer (TNBC) patient derived xenograft sample. Passages X1, X5, and X8 had 62, 369, and 231 cells post quality filtering (Methods section 9.1) respectively. We also include 17 mostly diploid control cells. These cells are combined to generate the input to the analysis pipeline (**Supplementary Fig. 2**). The second dataset, labelled *OV A* [1], consists of cells from three samples taken from a patient with high grade serous (HGS) ovarian cancer. The first sample, *SA*1090, was from an ascites pre-treatment, while *SA*922 was from an ascites post-treatment. The third sample, *SA*921, was taken from the ovary. See **Supplementary Fig. 3** for the tree and the CNA profile heatmap for this dataset. The final dataset, *SA*501 [1], is another TNBC xenograft tumour from 6 untreated passages, namely X2, X5, X6, X8, X11, and X15. After filtering, 515, 236, 328, 189, 836, and 308 cells remain in each passage respectively (for a total of 2,412 cells, see **Supplementary Fig. 4**). **Supplementary Table** 1 shows the attrition after each step of filtering cells per passage in each dataset.

To evaluate inferred trees from sitka and other tree reconstruction methods, we use a goodness of fit performance metric, which compares the compatibility of observed CN change points with a given phylogeny using Youden’s J index (Methods section 9.6, **Fig. 4-d**). Sitka has the highest Youden’s index across all three datasets. `UPGMA` and `WPGMA` perform similarly on *SA*501 and *SA*535. `UPGMA` performs slightly better than `WPGMA` on the *OV A* dataset. `HDBSCAN` has a close but slightly smaller Youden’s index than `UPGMA` over the *SA*535 and *OV A* datasets, but performs marginally better on *SA*501. `NJ` trails `WPGMA` on *SA*501 and the *OV A* datasets, and has the lowest Youden’s index on *SA*535. `MrBayes` performs well on the smallest dataset, *SA*535, with `MrBayes-np2` and `MrBayes-np8` performing similar to `WPGMA`, and `MrBayesWithBinaryInput` having achieved the second highest Youden’s index. On the *OV A* data, `MrBayesWithBinaryInput` and `MrBayes-np2` trail behind `medicc2` and `MEDALT`, while `MEDALTWithBinaryInput` has the lowest Youden’s index among all methods on all datasets. `MrBayesWithBinaryInput` and `MrBayes-np2` trail behind `NJ` over the *SA*501 dataset. `medicc2` and `MEDALT` without binary input, ran with default settings, did not yield a result given our available computational budget, with the former not finishing after several days, and the latter running out of memory (144 GB). Following [30], we run `MrBayes` for 10,000,000 generations. `MrBayes-np8` had completed only 278,000 iterations running on *SA*501 after several days. The results in this comparison suggest that sitka performs better than the baseline methods. While due to limitation to our available computational budget, we could not allocate more time to the benchmarked methods, it is possible that given more runtime/computation budget, the other methods might have converged to more accurate solutions. Running sitka on the real-world datasets took on average 22.3, 46.6, and 12.9 hours for the *OV A, SA*501, and *SA*535 datasets respectively, on a Linux workstation with 72 Intel Xeon Platinum 8272CL 2.60GHz CPU processors and 144 GB of memory.

### 2.3 Single cell resolution phylogenetic inference in PDX

Here we analyse the foregoing three multi-sample datasets. To visualise the tree inference results we arrange the inferred consensus tree *t* (Methods section 9.4.5) and the cell-by-bin CN matrix side by side where the rows of the matrix correspond to the position of individual cells on the tree and the markers are arranged by their genomic position (**Fig. 3-h**). **Fig. 4-a-c** shows examples of the multi-channel visualisation where each marker is represented by a tuple of three different data-types or *channels*, namely: (i) the latent markers induced by the consensus tree, *x*(*t*); (ii) the matrix of marginal posterior probability that cell *c* is a descendent of marker *l*, computed via the average 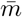 (**Fig. 3-g**, Methods section 9.4.5); and (iii) the sitka transformed input data *y*_*c,l*_.

We use this view to assess potential discrepancies between the input data and the inferred tree. In most cells and markers (as quantified in **Supplementary Fig. 9.6**), the observed data is in close agreement with the inferred tree. In the following we provide some examples of disagreements. Consider first the ChrX in the *OV* 2295 dataset (**Fig. 4-a**). ChrX has a long orange band (inferred marker in channel (i)) not matched by a black band (observed marker in channel (iii)) suggesting that a perfect phylogeny violation has occurred. The pattern in this marker is consistent with the presence of an ancestral event followed by a deletion. In **Fig. 4-b**, a set of diploid cells are attached to the root of the tree. These are control cells included in the experiment and correspond to a region in the bottom of the matrix with no inferred markers (orange bands) and almost no observed markers (black bands). In this dataset, there are change points where the observed marker has a high density (black band), but the tree is reconstructed with the marker absent (no matching orange band). Examples can be found in Chr1, Chr7 and Chr16. One possible explanation could be that the end-points of each event were detected as slightly shifted across cells. For instance, in **Supplementary Fig. 4** there are two bins with an amplification (CN state equal to three) in Chr1p where cells that harbour a mutation in the first bin appear not to have a mutation in the second bin, suggesting that the same event was called in the first bin in some cells, and in the second bin in others. An alternative hypothesis is that the cells in this dataset have a mutator phenotype that promotes CN mutations in these bins.

**Supplementary Fig. 5** shows the distribution of mismatch rates for each dataset, defined as the fraction of times that the observed and inferred markers do not match, i.e., 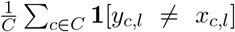 for *l* ∈ *L* (corresponding to the black and orange bands in **Fig. 4-a**). In *OV* 2295, 41 markers (11%) have a mismatch rate of over 50%, where marker *chr15_67000001_67500000* has the highest mismatch rate at 70%. In SA501, 30 markers (11%) have a mismatch rate of over 50%, 13 of which (5%) have a mismatch rate of over 75%. SA535 has the lowest maximum mismatch rate at 49% (marker *15_72000001_72500000*).

### 2.4 Placement of SNVs using the CNA inferred tree

To determine the presence or absence of SNVs in cells using data with high levels of missingness, we develop an extension of sitka, the sitka-snv model. Given single cell level variant read counts, the model incorporates CN data to place SNVs on the sitka-inferred phylogenetic tree. This *backbone* CN tree provides a principled way to pool statistical strength across groups of single cells sequenced at low coverage, including data from the DLP+ platform [16]. The output of the sitka-snv model is an *extended* tree that has marker nodes that comprise SNVs in addition to the original CNAs.

The SNVs are added to the existing CNA-based tree with the computational complexity of *O*(|*C*| + |*L*|) per SNV. **Fig. 7-c** shows the result of SNV placement with the number of variant reads in *SA*535, corresponding to the tree shown in **Fig. 4-c. Supplementary Figs. 14, 15**, and **16** show the number of variant reads and the matching SNV call probabilities for the *SA*535, *OV A* and *SA*501 datasets respectively. Sitka and sitka-snv provide a comprehensive genomic analysis tool for large scale low-coverage scWGS.

## 3 Discussion

Our method ignores certain pairwise dependencies induced by copy number change events having two end-points (except in cases where one of the end points is the end of the chromosome, e.g., whole-chromosome-arm events). This artificial duplication of the events having two input end-points can lead to the method being overconfident, i.e., outputting credible intervals that are smaller than expected. This is partly a reason for focusing more on point estimates (consensus trees) in this work, which we expect are less affected by this phenomenon (see Section 9.5.2).

In the present study we use data in which the genome of the single cells CNA profiles are partitioned into bins of a fixed size (500Kb), each assigned a constant integer CN state. The relatively large size is due to the low coverage inherent to the scWGS platform, but it implies that the same bin may harbour multiple CNA events. Biological processes that result in complex DNA rearrangements could further increase the probability of having two hits in one bin [31, 32]. Post hoc inspection is necessary to rule out large violations of our assumptions. This highlights the importance of our goodness-of-fit and visualisation methods as they help detecting such violations.

We note that we lose information when applying the sitka transformation. This transformation is necessary for the computational feasibility of the likelihood. In absence of this relaxation, the computational complexity of each iteration of the MCMC algorithm may no longer be bounded by *O*(|*C*| + |*L*|). Indeed, the approach to efficiently compute the likelihood depends on binary latent variables with specific perfect phylogeny assumptions, and it is not clear how to generalize this calculation to models that keep track of the evolution of more detailed CN state information along the phylogeny.

Structural variations such as chromothripsis, that affect multiple segments of the genome at the same time, make it difficult to determine the rate of CNA events and suggest that CNA events may not be suitable molecular clocks to estimate branch lengths. One possible remedy is to first infer the tree topology via markers based on CNA events and then conditioned on this topology, add SNVs to the tree. The number of SNVs on each edge of the tree may be used to inform branch lengths.

Our preprocessing pipeline excludes multiple cells from the analysis (see **Supplementary Table** 1). We filter out a fraction of cells to remove contaminated cells, either doublets (DNA material from two cells that is inadvertently merged) or mouse cells (in our real world datasets, we study human tumours that were transplanted into mice), cells with too many erroneous sequencing artefacts, and cycling cells (in the process of replicating their DNA). Removing a portion of the sequenced cells will decrease the statistical power to determine the subclonal structure of the population—an important application of this work—, and may bias the sampling against clones that have a higher division rate. We expect this will be an intrinsic limitation to any scWGS phylogenetic methods and this motivates the design of improved classification methods detecting cell cycling from genomic and imaging data.

We developed two main variants of our Bayesian models, one with error rate parameters shared by all loci (global) and one with locus-specific (local) error rates. In our simulation experiments, the global and local parameterizations performed similarly. Based on the similar performance of these two models and the fact that the global parameterization is computationally cheaper, we recommend the use of the global parameterization by default.

Evaluating the performance of a phylogenetic reconstruction method on real-world datasets is difficult, mainly due to a paucity of ground truth. One promising area of research is the use of CRISPR-Cas9 based lineage tracing [6]. In absence of ground truth data, we developed a goodness-of-fit framework that to our knowledge enables a first of a kind benchmarking of phylogenetic inference methods over real-world scWGS CNA datasets.

Phylogenetic tree reconstruction is a principled way to identify subpopulations in a hetero-geneous single-cell population. This in turn enables the use of population genetics models that track the abundance of subpopulations over multiple timepoints [5] and to make inferences about the evolutionary forces acting on each clone. Further study with timeseries modelling will provide insight into therapeutic strategies promoting early intervention, drug combinations and evolution-aware approaches to clinical management.

## 4 Acknowledgements

This project was generously supported by the BC Cancer Foundation at BC Cancer and Cycle for Survival supporting Memorial Sloan Kettering Cancer Center. SPS holds the Nicholls Biondi Chair in Computational Oncology and is a Susan G. Komen Scholar (#GC233085). SA holds the Nan and Lorraine Robertson Chair in Breast Cancer and is a Canada Research Chair in Molecular Oncology (950-230610). Additional funding provided by the Terry Fox Research Institute Grant 1082, Canadian Cancer Society Research Institute Impact program Grant 705617, CIHR Grant FDN-148429, Breast Cancer Research Foundation award (BCRF-18-180, BCRF-19-180 and BCRF-20-180), MSK Cancer Center Support Grant/Core Grant (P30 CA008748), National Institutes of Health Grant (1RM1 HG011014-01), CCSRI Grant (#705636), the Cancer Research UK Grand Challenge Program, Canada Foundation for Innovation (40044) to SA, SPS and ABC. We extend our gratitude to Sarah P. Otto for her helpful comments on a draft of this manuscript.

## 6 Author Contributions

SS, FD: computational method development, data analysis, manuscript writing; KC: data analysis, manuscript writing; KRC, AR: method development; FK, data generation; DL, MA, AM, MW, TF: computational biology, data analysis; NR: manuscript editing; SA: data generation and oversight; SPS: method development and oversight; ABC: project conception and oversight, statistical inference method development, manuscript writing, senior responsible author.

## 8 Code availability

Sitka is available at https://github.com/UBC-Stat-ML/sitkatree.git.

## 9 Methods

### 9.1 Pre-processing

The raw data contain cells that are either contaminated (e.g., contains biological material from mice) or have undesired sequencing artefacts. These include cells that were captured for DNA sequencing when undergoing mitosis. Since the sitka model does not account for such phenomena, the filtering is an important step. **Supplementary Fig. 12** shows the steps taken from pulling the raw data to the CNA integer matrix ready for sitka transformation (details in the Supplementary Information). Briefly, we remove control cells, cells with highly-noisy CN calls, and cells that have very few mapped reads. We also remove copy number bins that lie in difficult to sequence regions of the genome (bins with lowmappability). Finally, we drop cells that, based on their CNA profile, are suspected to be cycling cells.

### 9.2 The sitka transformation

To obtain the *C* × *L*_Markers_ phylogenetic markers matrix *y* that comprises the input to the sitka model, we apply a lossy transformation to the *C* × *L*_Bins_ CNA matrix *a* that involves computing the change in copy number state between two consecutive bins. **Supplementary Fig. 1** shows a small CNA matrix and its corresponding transformation into the marker matrix. For brevity, in what follows we assume that only one chromosome is used, so that *L*_Bins_ = *L* and *L*_Markers_ = *L*_Bins_ − 1. In practice, we use all available chromosomes, and *L*_Markers_ = *L*_Bins_ − *N*_Chr_ where *N*_Chr_ denotes the total number of chromosomes used.

Given a filtered cell-by-locus matrix *a*, we sort bins by their genomic position. Then in each chromosome, we compute markers as the binarised difference between consecutive bins. In other words, *y* = (*y*_*c,l*′_) and *l*′ ∈ {1, …, *L* − 1}, and

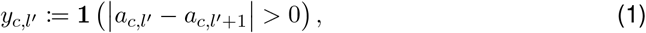

where **1**(*x*) is the indicator function.

### 9.3 Fixing jitter and selection of phylogenetic markers

The copy numbers available to us in this work are estimated independently for each cell. This is one reason why the start position (bin) of the same CN change event may be slightly different across cells, generating some *jitter*. We address this by enumerating each change point column in order of decreasing density (where the density of column *l* is given by Σ_*c*∈*C*_*y*_*c,l*_/|*C*|) and merging the column with its *k* = 2 immediate neighbours (see Algorithm 1 for details). An example of the result of the jitter correction heuristic is shown in **Fig. 3** panel **c**. To speed-up computation, only a subset of markers present in at least a minimum number of cells are chosen for phylogenetic inference. That is, we removed columns *l* in *y* with relative density Σ_*c*∈*C*_*y*_*c,l*_/|*C*| less than a threshold, set to 5%. Larger values of this threshold may lead to less resolved clades in the inferred tree.

#### Algorithm 1 JitterFix

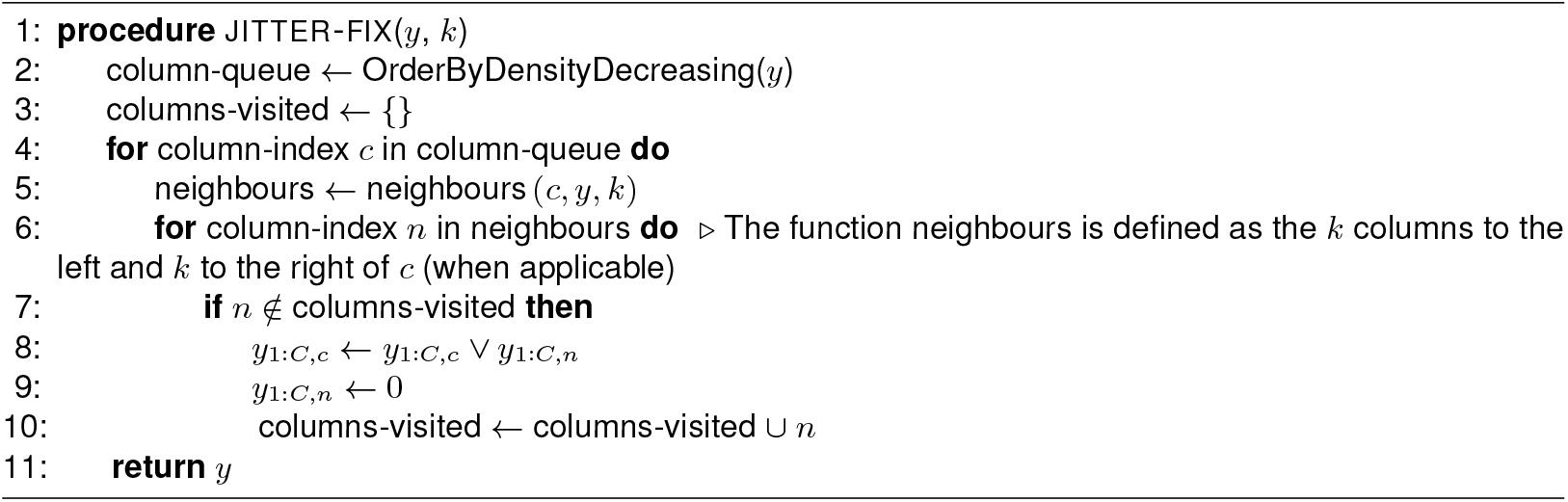

### 9.4 The sitka model

#### 9.4.1 Model description

The sitka model starts with the perfect phylogeny assumption for the latent variables *x*_*c,l*_ but allows deviation from it via allowing noisy observations *y*_*c,l*_. In a perfect phylogeny model, each phylogenetic trait arises only once on the rooted tree topology and all cells descending from that position will inherit that trait and no deletions are allowed.

Let *C* and *L* denote the sets of cells and loci respectively.

We posit an observation probability model *p*(*y*|*x, θ*), where *θ* are model parameters described shortly, and both *x* and *y* are cell by locus matrices, the former being latent (derived from the unobserved tree via *x* = *x*(*t*)), while the latter is the matrix obtained from the sitka transformation. To model errors in copy number calls as well as perfect phylogeny violations, we introduce false positive and negative rate parameters *r*^FP^ ∈ (0, 1) and *r*^FN^ ∈ (0, 1) respectively, and an error matrix

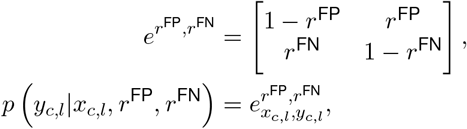

from which we set:

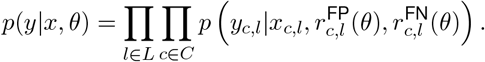

We define two type of models, differing in the choice of functions 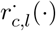 and dimensionality of *θ*: one based on global error parameters, and one based on locus-specific error parameters.

For the global parameterization, 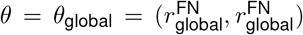, and the false positive and false negative functions are given by 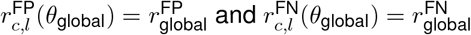.

For the locus-specific error model, we set the error rates to be locus-dependent: 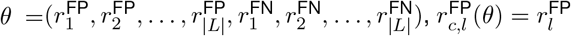 and 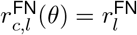. With this extra flexibility, the model can discount the effect of a trait violating the perfect phylogeny assumption, by setting high error rates for the trait’s locus.

The two parameterizations are compared in the Supplementary Information. We use the global parameterization by default unless mentioned otherwise.

In both the global and locus-specific parameterizations, we need to construct a prior distribution *p*(*θ*) over the error parameters. Using a uniform prior distribution with support on [0, 1] can lead to pathological cases as shown in **Fig. 2**. To avoid that, we use the following prior distributions on the two types of error:

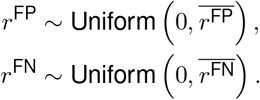

We use 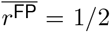 and 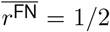 in our experiments involving synthetic data. For experiments on real world data, we use 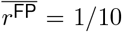 and 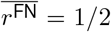 as default. When the model is misspecified from an overly conservative bound, the trace, and thus posterior distribution, collapses to the boundaries. For example, when using a false positive rate of 0.1 for synthetic data, the resulting approximate marginal posterior of the false positive rate corresponds to a near-point mass at 0.1. We did not observe such boundary collapse on the real datasets studied in this work.

Next, we describe the prior *p*(*t*) on phylogenies using a two-step generative process:

##### Sampling a mutation tree

let 𝒱^m^ = *L* ∪ {*v*\*}denote a vertex set composed of one vertex for each of the |*L*| loci plus one artificial root node *v*\*. The artificial root node induces an implicit notion of direction on the edges, viewing them as pointing away from *v*\*. Let 𝒯^m^ denote the set of trees *t*^m^ spanning 𝒱^m^. The interpretation of *t*^m^ is as follows: there is a directed path from vertex/locus *l* to *l*′ in *t*^m^ if and only if the trait indexed by *l* is hypothesized to have emerged in a cell which is ancestral to the cell in which *l*′ emerged. Pick one element *t*^m^ ∈ 𝒯^m^.

##### Sampling cell assignments

assign each cell to a vertex in *t*^m^. The interpretation of assigning cell *c* to locus *l* is that among the traits under study, *c* is hypothesized to possess only the traits visited by the shortest path from *v*\* to *l* in *t*_*m*_. If a cell *c* is assigned to *v*\*, the interpretation is that *c* is hypothesized to possess none of the traits under study.

The number of possible trees obtained from this two-step sampling process is:

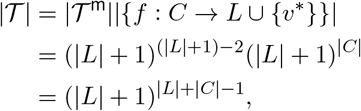

where we use Cayley’s formula to compute |𝒯^m^|. Hence the uniform prior probability mass function over the possible outputs of this two-step sampling process is given by:

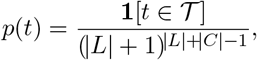

where 𝒯 is the set of all perfect phylogenetic trees that result from the two step generative process described above. Simulation from the prior can be performed using Wilson’s algorithm [33], followed by independent categorical sampling to simulate the cell assignments.

This simple prior has a useful property: if a collection of say two splits are supported by *m*_1_ and *m*_2_ traits, then the prior probability for an additional trait to support the first versus second split is proportional to (*m*_1_ + 1, *m*_2_ + 1). Therefore, there is a “rich gets richer” behaviour built-in into the prior, which is viewed as useful in many Bayesian non-parametric models [34]. More precisely, this “rich gets richer” behaviour emerges when grouping trees into equivalence classes and looking at the induced prior on these equivalence classes, i.e., the distribution obtained by summing the prior over the trees in the equivalence class. Specifically, consider the equivalence relation such that two type I trees *t, t*′ are in the same equivalence class if and only if *f* (*t*) = *f* (*t*′), where *f* (·) consists in transforming *t* into a type II tree while annotating each edge by the number of events on that edge. Since there are different numbers of type I trees in different equivalent classes, this means that the induced prior on these equivalence classes is non-uniform.

#### 9.4.2 Inference

The posterior distribution,

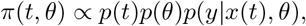

is approximated using MCMC. Two MCMC moves are used, described in the next two sections. The posterior distribution is summarized using a Bayes estimator described in Section 9.4.5. The model is implemented in the Blang probabilistic programming language [35].

#### 9.4.3 MCMC tree exploration move

Sitka uses a tree sampling move to efficiently explore, at each MCMC iteration, the posterior distribution in a large neighbourhood of a given tree. Given a tree *t* and locus *l*, we define a neighbourhood *N*^*l*^(*t*) ⊂ 𝒯 by removing *l* from *t*, and considering all possible ways to reattach *l* and hence defining a neighbourhood of phylogenetic trees (we also implemented a separate move reattaching cell nodes instead of locus nodes, its derivation follows similar lines as the move described in this section). The process of removing *l* is called an *edge-contraction* (removing an edge after connecting its two end-points) while the process of adding back a locus is called an *edge-insertion*. An edge insertion (see **Supplementary Fig. 1** for a visualization) can be described as follows:

1. Pick a non-cell vertex *v*, i.e., an element from the set *R* = {*v*\*} ∪ *L*\{*l*} where *v*\* is the root node.
2. Pick any subset of *v*’s descendent subtrees and disconnect them from *v*.
3. Add a new node *l* under *v* and move the selected nodes from step 2 above and attach them to *l*.

In the following, we derive the probability distributions to be used in steps 1 and 2 above that lead to a Gibbs sampling algorithm [36]. The Gibbs sampler first selects a locus *l* from a fixed distribution (a tuning parameter), which we take for simplicity as being uniform over the |*L*| loci.

After having sampled *l*, we partition *N*^*l*^(*t*_\*l*_) into blocks corresponding to the choice of node *v* made in Step 1, 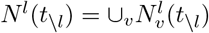. The Gibbs conditional probabilities required in step 1 above are of the form:

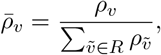

where:

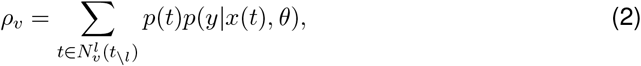

and *t*_\*l*_ denotes the tree obtained after performing an edge contraction, where the contracted edge is between *l* and the parent node of *l*. To compute *ρ*_*v*_ efficiently, we start with the following likelihood recursion for all vertex *v* in *t*_\*l*_. First, for all vertices *c* corresponding to a cell and *b* ∈ {0, 1}, define:

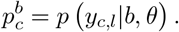

Next, we perform the following bottom-up recursion for all subtrees of *t*_\*l*_ : for all *v* ∈ *R, b* ∈ {0,1},

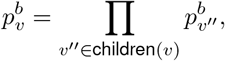

where children(*v*) denotes the list of children of vertex *v*.

We can now return to the problem of computing 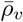. First, observe that the sum in Equation (2) can be re-indexed by a bit vector ***b*** = (*b*_1_, *b*_2_, …, *b*_*k*_), *b*_*v*″_ ∈ {0, 1} of length equal to *k* = |children(*v*) |. Each bit *b*_*v*″_ is equal to one if children *v*″ is to be moved into a child of *v*′ (refer to **Supplementary Fig. 1**), and zero if it is to stay as a child of *v*. For each possible assignment, we obtain a tree 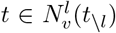, and its probability can be decomposed into factors corresponding to cells that are descendant of *v* (denoted *C*_*v*_, solid red thick line under the tree of **Supplementary Fig. 1-B**) and those that are not (denoted *C*_\*v*_, dashed green thick line under the tree of **Supplementary Fig. 1-B**).

The product of the likelihood factors corresponding to cells that are not descendants of *v* (“outside product”) does not depend on the choice of the bit vector. This outside product can be obtained as follows:

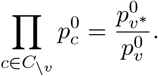

Note that this assumes 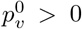. As a workaround to cases where there are structural zeros, we recommend injecting small numerical values if 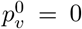 (we used 10^−6^ in our implementation).

For the cells under *v*, we now have to take into account whether they are selected under the newly introduced locus or not. More precisely, for each of the children *v*_1_, *v*_2_, …, *v*_*k*_, we have to take into account the value of the bit vector *b* = (*b*_1_, *b*_2_, …, *b*_*k*_). The sum over possible assignments written naively has a number of terms which is exponential in *k*, but can be rewritten into a product over *k* factors:

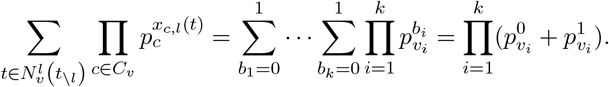

Putting it all together, we obtain for some constants *K*_*i*_ independent of *v*:

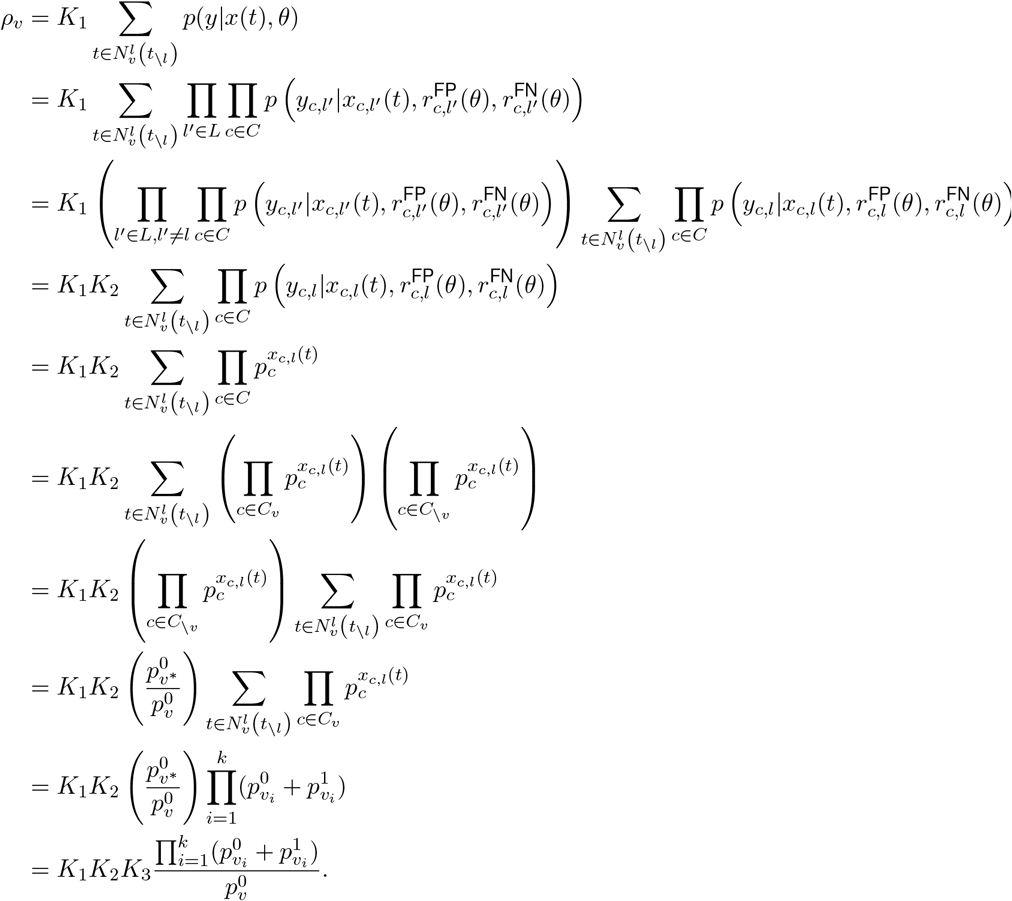

Putting these together we can compute the probabilities required in step 1 above:

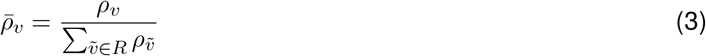

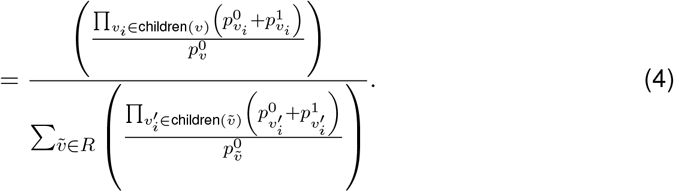

Once *v* is sampled, we choose a subset of its children to move to *v*′ by sampling *k* independent Bernoulli random variables with the *i*-th one having bias

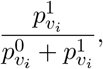

and selecting children with corresponding Bernoulli realisations of 1.

#### 9.4.4 MCMC parameter exploration move

To resample the parameters *θ* we condition on the tree *t*, and hence on the hidden state matrix *x*, and update *θ* in a Metropolis-within-Gibbs framework. There are two different samplers depending on whether the global or locus-specific parameterization is used. We start with describing the former.

We compute two sufficient statistics from the matrix *x* (i) the number of false positive instances, *n*^FP^, and (ii) the number of false negative instances, *n*^FN^,

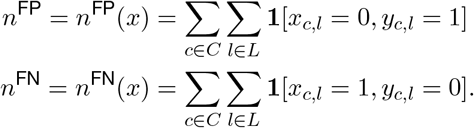

Based on these cached statistics, we obtain:

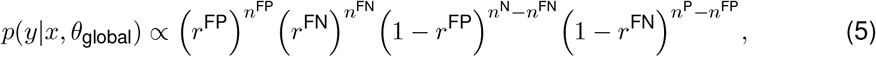

where the the number of positive *n*^P^ and negative *n*^N^ instances in the data can be pre-computed,

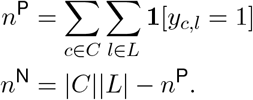

Based on the above expression, which can be evaluated in *O*(1) once the statistics are computed, we then use a slice sampling algorithm to update the parameters [37].

The sampler for the locus-specific parameterization is very similar. The main difference is that we compute the statistics for each locus *l*:

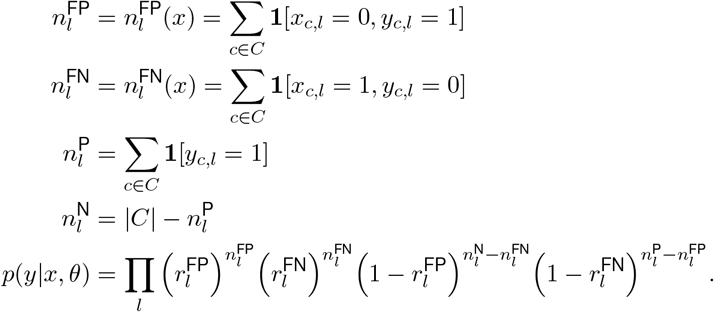

Then a slice sampling move is applied to each locus-specific parameter.

#### 9.4.5 Posterior summarization

To summarize the posterior distribution using a point estimate, we approximate the Bayes estimator [38] by minimising the Bayes risk for a loss function 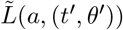 encoding the cost of selecting an “action” *a* when the true tree is *t*′ and the true parameter is *θ*′:

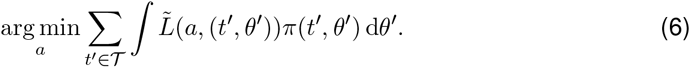

Here, an “action” consists in selecting a consensus tree *t*. Moreover, the loss function we consider only depends on the true tree *t*′ and not on the true parameter *θ*′, so we write this loss function as *L*(*t, t*′), simplifying the above equation into:

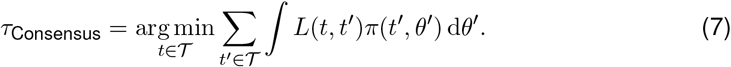

One default choice for *L*(*t, t*′) is the *L*_1_ metric on the matrices of induced indicators *x*(*t*), *x*(*t*′):

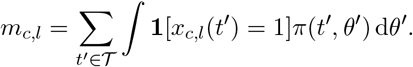

It is useful to define the marginal indicators *m*_*c,l*_ that can be conceptualised as the posterior probability of cell *c* to have trait *l*:

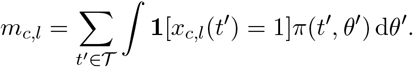

Using the MCMC samples *t*^1^, *t*^2^, …, *t*^*N*^, we obtain a Monte Carlo approximation:

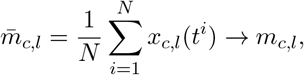

with probability one.

**Fig. 3-g** shows an example of the matrix *m* each element of which is one of the approximated 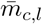. We can now write the objective function of Equation (7) via the above marginal indicators:

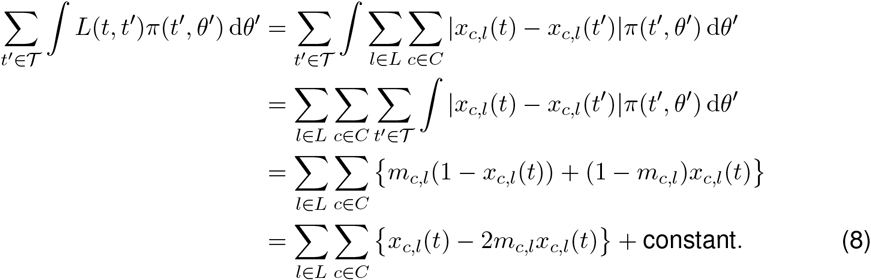

We use a greedy algorithm to approximately minimize Equation (8). We start with a star tree with leaves *C* rooted at *v*\* and add loci from *L* one by one from a locus queue sorted by priority score. The priority score of each locus *l* is computed as

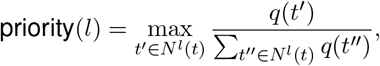

where

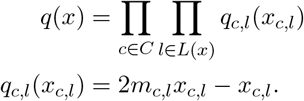

The quantities in the priority queue can be computed as in Section 9.4.3. We take the result of the minimization of the Bayes risk as the consensus tree *τ*_Consensus_.

#### 9.4.6 Consensus tree and CNA heatmap visualisation

To visualize the consensus tree, we collapse the chains (sequence of loci having only one child) as well as remove the subtrees containing no cells. We align the leaves of the tree which correspond to cells after collapsing to the rows of a cell-locus matrix.

### 9.5 Synthetic experiments

#### 9.5.1 Benchmarking

To assess the performance of sitka against alternative approaches, we ran inference on 90 simulated datasets of varying characteristics. We will refer to this set of datasets as *S*90; its simulation procedure is described in Section 9.5.3. For each dataset in *S*90, we scored each method by computing the Robinson-Foulds (RF) [39] distance between the simulated tree and the inferred tree. The scores were normalized within each dataset by dividing each method’s score by the worst performing method’s score (note that the set of methods includes sampling a tree uniformly at random; the motivation of this normalization is to correct for the intrinsic difficulty of datasets).

We compared sitka against the following baseline methods: `UPGMA`, `WPGMA`, `NJ`, `HDBSCAN`, and balanced and ordinary least-squares minimum-evolution methods (`BME`, `OME` respectively) of [40]. We also report the score of a uniformly random bifurcating tree, `Uniform`, to help interpret the absolute scores. Each method was given raw data from *S*90, as well as input identical to that of sitka, i.e., filtered binary marker data. Sitka’s inference settings are summarized in **Supplementary Table 2**.

Baseline methods performed significantly worse with sitka’s input and are thus omitted from the following summary. Sitka’s normalized RF score (0.57 ± 0.04) outperformed all baseline methods, the next best performer was `BME` (0.91 ± 0.07). Sitka ranked first in all 90 but one set of data, where it ranked third for one dataset of size 500 × 800. These results are summarized in **Supplementary Fig. 8**.

#### 9.5.2 Exploratory experiments within sitka

To explore the effectiveness of global versus *local* (locus-specific) parameterization (Section 9.4.1), and the posterior summarization method (Section 9.4.5), we ran inference on 10 synthetic datasets. We will refer to this set of datasets as *S*10; its simulation procedure is described in Section 9.5.3. Inference settings are summarized in **Supplementary Table 2**.

RF distances from the *best-possible tree* were computed as a metric. The best-possible tree is defined as the perfect phylogenetic tree constructed from the noiseless synthetic, unviolated cell-locus matrix data. Note that the best possible tree is derived from the *true tree* (i.e., the one used in the first step of the generating process), but in general can be different. To understand why, note that the tree generation process will simulate on each edge of the true tree a Poisson distributed number of evolutionary events (Section 9.5.3). As a result, some edges can have zero associated evolutionary events. This means that even if we turned off all observation noise it would not be possible to recover these zero-event edges, they are in a sense unidentifiable. The process of producing the best possible tree essentially consists in collapsing these zero-event edges, hence forming a multifurcating reference tree.

For a baseline with which to compare the greedy estimator (GE) of Section 9.4.5, consider the *trace search estimator* (TSE). The TSE is defined as a tree in the sampler trace that minimizes the sample *L*_1_ distance (Section 9.4.5). Formally,

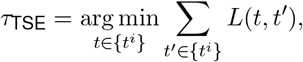

where {*t*^*i*^} denotes the set of trees that were sampled during the MCMC procedure.

The GE method outperformed the TSE method under both the global and local models. This suggests the proposed GE can, informally, harness more information from the posterior. Under the TSE, the global model (0.44 ± 0.09; mean normalized RF score ± standard error) outperformed the local model (0.71 ± 0.06). This observation suggests that the local parameterization has a strong influence on the trace (in tree space) of our sampler, as the TSE is essentially a search over the posterior sample. Under the GE, the global model (0.31 ± 0.07) and local model (0.30 ± 0.07) performed evenly well. This observation suggests that the choice of parameterization does not heavily influence the information contained in the marginal posterior over trees. Ultimately this experiment suggests that the GE summarizes the marginal posterior sufficiently well such that the global model, the simpler model of the two, suffices for reconstructing phylogenies and should be the preferred model. A summarizing plot is shown in **Supplementary Fig. 9**.

In our next synthetic experiment, we aimed to study the effects of perfect phylogeny assumption violations on the reconstruction of trees, and attempted to draw connections to real world data. The two violations considered are infinite sites and loss violations, described in Section 9.5.3. Inference was performed on 130 datasets (*S*130). Inference settings are summarized in **Supplementary Table 2**, and the simulation procedure for *S*130 is described in Section 9.5.3.

The experiment results are summarized in **Fig. 7-a**. Holding one violation rate fixed at zero and varying the other, we observed linear effects for both types of violations. The results suggest sitka is more robust to infinite sites violations, with estimated effects to be 0.31 ± 0.07 (normalized RF distance ± standard error), which is much less than loss violations (0.47 ± 0.07). When varied together, the linear effects were estimated to be 0.25 ± 0.04, 0.38 ± 0.04 respectively. In an attempt to draw connections to real datasets, we developed a heuristic method to obtain a rough estimate of both violation rates. The estimated rates obtained on real data were all less than 0.25 (the estimation heuristic is described below; **Fig. 7-b**).

We now describe the heuristic we used to obtain rough estimates of the rates of the two types of violation. Given the inferred tree and its corresponding marker matrix *x* (as in Section 9.4.1), and the sitka-transformed marker matrix *y* (as in Section 9.2), define the difference matrix *z* := *x* − *y*, i.e., *z* has entries *z*_*i,j*_ = *x*_*i,j*_(*t*) − *y*_*i,j*_, where *t* is the consensus reconstruction. To motivate how we can detect violations of the Infinite Site assumption (IS, i.e., genomic bins in which more than one events occur), and losses, refer to 10, and notice that these violations tend to leave a distinctive pattern on the difference of the two matrices *x* (positive entries shown in black in the Figure) and *y* (positive entries shown in orange in the Figure). Specifically, define *z*_Loss_ with entries 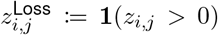, and similarly *z*_IS_ with entries 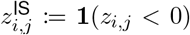. Given an integer-valued threshold *ϵ*_v_*>* 0, we say a column or trait *l* in *z*_*v*_ (for *v* ∈ {Loss, IS}) has a violation if there exists an *island* of size at least as large as *ϵ*_v_. An island of size *s* in column *l* is defined to be any sequence of row indices *i, i* + 1, …, *i* + *s* such that 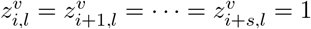 and 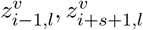 are, not necessarily the same, 0 or undefined. Finally, the proportion of columns with a given type of violation, loss or infinite sites, is taken to be the violation rate estimate.

We also performed experiments to compare our full Bayesian analysis to Maximum Likelihood estimation (MLE). To do so, we generated data as follows: we first sampled a tree generated uniformly over topologies; second, we generated synthetic data according to *y*_*c,l*_|*x*_*c,l*_ ∼ Normal(*x*_*c,l*_, *σ*^2^), varying *σ*^2^ to control the amount of noise. We generate matrices of size |*C* |= 1000 and |*L*| = 50 for *σ*^2^ ∈ {1/10, 2/10, …, 5/10}. In these experiments we provide the well-specified noise model to both inference methods. We approximate the MLE using a greedy scheme with the same structure as the one described in Section 9.4.5. The results are shown in Figure 8. The Bayesian methods outperform the greedy maximum likelihood heuristic by a large margin.

**Figure 8.**
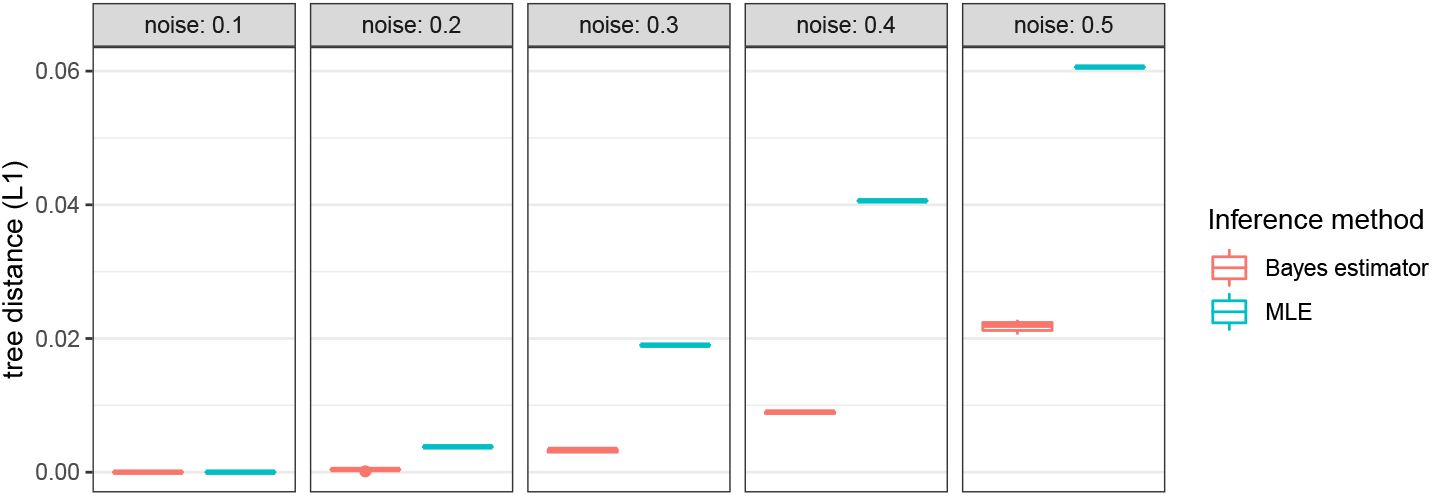
Synthetic data results comparing our Bayesian estimator to a Maximum Likelihood Estimator (MLE). Boxes from left to right show different amounts of noise in the synthetic data generation, corresponding to values for *σ*^2^. The *y* axis measures the L1 tree distances loss, normalized by |*C*||*L*|.

Finally, we investigate the impact of ignoring pairwise dependencies between the two end points of CNA events. We first make the observation that if we subset the sitka markers to keep only those where the copy number is increasing from left to right, we retain only one end point of each paired event. This creates a smaller set of independent markers *L*′ *⊂ L*. We can compute one sitka tree *t* based on all *L* loci (which includes ignored pairwise dependencies), and one sitka tree *t*′ based on *L*′ (a smaller set of independent loci). We can then inspect the proportion of identical entries in the matrices *x*(*t*′) compared to *x*(*t*), the latter subsetted to the columns in *L*′.

We performed the experiment described above on the S90 datasets (described in 9.5.3) with three noise regimes described as follows: (I) where step (ii) in 9.5.3 is skipped; (II) uniform noise parameters FPR and FNR drawn from uniform distributions on the intervals (0.0005, 0.005), (0.005, 0.015) respectively, doubling noise parameters drawn from a uniform distribution on (0.015, 0.035) distribution, jitter noise parameters drawn from a uniform distribution on (0.15, 0.35); (III) uniform noise parameters FPR and FNR drawn from uniform distributions on the intervals (0.001, 0.01), (0.01, 0.03) respectively, doubling noise parameters drawn from a uniform distribution on (0.03, 0.07) distribution, jitter noise parameters drawn from a uniform distribution on (0.3, 0.7). All results are averaged over 15 datasets.

In all three noise regimes we observed a large overlap between *t* and *t*′, but this overlap is negatively correlated with noise: in regime (I) we observed a mean overlap of 0.99 (sd 0.004); in regime (II), a mean overlap of 0.97 (sd 0.009); in regime (III), a mean overlap of 0.76 (sd 0.18). The results support that in a low to moderate noise regime, it is reasonable to ignore violation of pairwise dependencies for the purpose of point estimation (consensus tree construction). In the higher noise regime, it may be advantageous to build the two trees *t* and *t*′. We expect neither to systematically outperform the other, the trade-off being that *t* is built from more data but with independence violations, whereas *t*′ is built from less data but without independence assumption violations. Our goodness-of-fit tests can be used to select one of these two trees for final output.

#### 9.5.3 Data simulation

Datasets in *S*90 were generated in two steps: (i) simulate a cell tree and its corresponding CNA data, and (ii) inject noise into the CNA data from step one.

In the first step we used the simulator of [41] to generate trees along with CNAs, where leaf nodes represent observed cells and internal nodes represent latent ancestral cells, i.e., unobserved cells. An edge in the tree represents an ancestral relationship between the respective cells.

The simulator of [41] itself consists of two parts, which we briefly describe as follows. First, the simulator samples a tree based on a generalization of the Blum-François Beta-splitting (GBFBS) model [42, 43], which is inspired by the Beta-splitting model of [44].^1^ The Beta-splitting model is particularly well-suited for generating a wide range of topologies, varying from balanced to imbalanced tree structures. Second, given a tree, CNAs are simulated on the edges of the tree where the number and size of CNAs are drawn from Poisson and exponential distributions respectively. The simulator also accounts for clonal whole chromosome amplification events, motivated by punctuated evolution models [45].

The second step of our synthetic data simulation process, independent of [41], injects noise into a cell by locus input CNA matrix *y*, and outputs a noisy matrix of the same size. Three types of noise were employed, namely, uniform noise, jitter noise, and a doubling noise.

The uniform noise is parameterized by false positive (FPR) and false negative (FNR) rate parameters. For each element of the input matrix *y*_*ij*_, add an integer *N*_*ij*_ ∼ Binomial(*y*_*ij*_, FNR) or subtract an integer *M*_*ij*_ ∼ Binomial(1, FPR).

The doubling noise is parameterized by a probability *p*_d_: for each row of the CNA matrix *y*, draw a factor *K* where *K* − 1 ∼ Binomial(1, *p*_d_), which is then multiplied to the row of the CNA matrix as noise. This procedure effectively, on average, doubles the copy number values for *p*_d_ proportion of cells in the sample.

The jitter noise is parameterized by a probability *p*_j_. First, map the CNA matrix to its marker matrix. Then for each marker, the locus corresponding to the marker is randomly duplicated to the previous bin(s), or the next bin(s). The number of bins *J* to be overwritten — zero, one, or two — is drawn from a Binomial(2, *p*_j_) distribution.

Datasets in *S*90 were of sizes {500, 1000, 1500, 2000, 2500, 3000} cells by (approximately) {400, 600, 800} markers. For each combination of sizes, we generated five datasets based on different random seeds and parameters to make a total of 6 × 3 × 5 = 90 datasets. The approximate number of markers is the target number of markers after correcting for jitter and filtering. **Supplementary Fig. 7** shows the CNA profiles of a subset of simulated data.

To describe the simulation parameters used for *S*90, we follow the terminologies and notation used in [41]. For generating trees, the *α* and *β* values parameterize the generalized Beta-splitting model. We used a symmetric parameterization of *α* = *β* ∈ {− 0.9, − 0.83, − 0.7, − 0.48, − 0.1}. For generating CNA data, the mean number of CNA to be added to a branch in the tree was chosen to generate data with approximately the number of desired markers post filtering and jitter-fixing. The multiplier of the mean CNA on the root was set to 8, the whole amplification rate (rate of an allele chosen to be amplified) was set to 0.5. The remaining parameters used default settings. See [41] for a more thorough description of parameters.

For injecting noise, we drew the uniform noise parameters FPR and FNR from uniform distributions on the intervals (0.001, 0.01), (0.01, 0.03) respectively. The doubling noise parameters *p*_d_ were drawn from a Uniform(0.03, 0.07) distribution. The jitter noise parameters *p*_j_ were drawn from a Uniform(0.3, 0.7) distribution.

Datasets in *S*10 and *S*130 were also generated in two steps: (i) simulate a cell tree and its corresponding binary marker data satisfying perfect phylogeny assumptions, and (ii) inject noise and/or violations into the the binary marker data from step one.

In the first step, a tree is generated via Kingman’s coalescent [46].^2^ Briefly, we sample a coalescent tree for the set of cells *C* by uniformly selecting pairs of cells *c*_*i*_, *c*_*j*_ ∈ *C* to coalesce backwards in time. The waiting time, or the branch length, between each event is exponentially distributed. Conditionally on the coalescent tree and given a set of loci *L*, we simulate a |*C*| × |*L*| marker matrix *y*. Every entry *y*_*i,j*_ is initialized to 0. Then for each column *l*, we select a subset of cells *C*′ from *C* to set *y*_*i,l*_ to 1, for all *i* ∈ *C*′. The subset of cells is sampled by choosing a branch on the tree with probability proportional to the branch length, and selecting all cells descendant from the selected branch. In essence, we are simulating the number of events via a Poisson process, and directly mapping these events to the cell-locus marker matrix. The above concludes the data generation procedure satisfying perfect phylogeny assumptions.

In the second step of *S*10’s simulator, we injected artificial noise by introducing standard false positive and negative values into *y*. This concludes *S*10’s simulator. The simulator for *S*130 has an additional sampling step for controlling the degree of perfect phylogeny violations. We considered two types of violations: (i) the loss of markers along a tree’s branches, and (ii) the violation of the infinite sites (IS) assumption, that is, the occurrence of multiple distinct events in the same locus.

The procedure for simulating loss of marker events can be described as follows. First, randomly select a locus *l*, then identify the most recent common ancestor *a* for the set of cells {*i* : *y*_*i,l*_ = 1}. Given *a*, sample a cell *d* descendant of *a* (including *a*). Finally, the loss event is simulated by reverting *y*_*i,l*_ to 0, for all *i* descendant of, and including, *d*.

IS model violations were simulated as follows. Uniformly sample a pair of loci (*j, k*), and merge *y*_·,*j*_, *y*_·,*k*_ into one column, yielding a cell-locus matrix of size one less than the original size. However, to maintain control over |*L*|, datasets in *S*130 were simulated with |*L*| + *N*_IS_ loci such that after simulating IS violations, we recover a matrix of size |*C* |×|*L*|, where *N*_IS_ is the number of IS violations.

The total number of loss and infinite sites violation events (*N*_Loss_, *N*_IS_) were drawn from binomial distributions with probability *p*_Loss_, *p*_IS_ respectively (and size |*L*|). As a final step, false positives and negatives were artificially injected.

For both *S*10 and *S*130, datasets of size |*C*| × |*L*| = 500 × 100 with FNR and FPR both set to 0.002 were generated. For *S*130, the unordered pair (*p*_Loss_, *p*_IS_) were set to values in {(0, 0), (0.1, 0.1), …, (0.4, 0.4)} *∪* {(0, 0.1), (0, 0.2), (0, 0.3), (0, 0.4)}. For each configuration of simulation parameters, 10 different seeds were used to generate a total of 10 and 130 datasets for *S*10 and *S*130 respectively.

### 9.6 Goodness-of-fit

To evaluate the goodness-of-fit of inferred trees on real data, we suggest a test comparing the posterior distribution over entries of the matrix *x* with the data *y*.

Since we will assess the goodness-of-fit of not only our method but also different baseline, we start by explaining how we can generalize the notion of the *x* matrix used in our method to other tree reconstruction methods. To do so, consider an inferred rooted tree, *τ*, and define a matrix-value function *g*(*τ*) as follows. If *τ* is a tree inferred from sitka, set *g*(*τ*) = *x*(*τ*). For trees inferred from baseline methods, we proceed as follows. Let *τ* denote a rooted tree, *u*, one of its unlabelled internal nodes, and *c* one of its leaves. Let clade(*u*) denote the clade corresponding to *u*, i.e., the set of leaves descendent from *u*. We define *g*_*c,u*_(*τ*) = **1**[*c* ∈ clade(*τ*)].

In general the inferred trees from the baseline methods do not have named internal nodes, nor do they have the same number of internal nodes as the number of loci *L*. Therefore we do not know which locus in the inferred tree *τ* corresponds to which locus in the matrix *y*. We note that this is not the case with trees inferred from sitka where the internal nodes of the tree correspond to the columns of the induced genotype matrix *g*. As a result, for methods other than sitka, for each column in the input data matrix, we pick a clade in *τ* that has the highest prediction accuracy for the entries in that column.

More precisely, for each method, we report Youden’s *J* index [49] which is equal to the sum of the sensitivity and specificity minus 1. We now define a binary classification counts matrix function *h*, i.e., a function which, for two vectors *w* and *z* of length *C*, forms the confusion matrix:

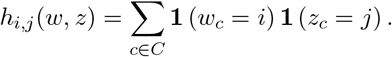

For example *h*_0,0_(*w, z*) would count the number of times both elements of *w* and *z* were equal to zero (or *true negative*). We define accuracy for a given confusion matrix *h* = *h*(*w, z*) as

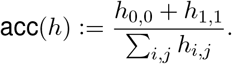

We further define sensitivity and specificity as

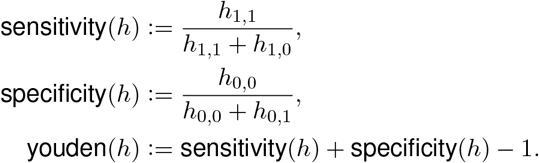

For a given tree *τ* and its corresponding genotype matrix *g* = *g*(*τ*) we compute the Youden’s score as follows:

1. for all locus *l* in *y, h*_*l*_ := arg max_*l*′∈columns(*g*)_ acc *h*(*y*_*l*_, *g*_·,*l*′_),
2. *h*_*τ*_ := Σ_*l*_ ′_∈columns(*g*)_*h*_*l*′_
3. youden_*τ*_ := youden(*h*_*τ*_).

That is for each locus in *y*, we take the clade that among all possible clades in *τ* maximizes the accuracy in predicting which cells are present in the *l*-th column of *y*. We then sum over all these scores to compute a confusion matrix for *τ* and use this agglomerative matrix to compute the Youden’s score for the tree. We use the delta method to calculate confidence intervals. Recall that the delta method is concerned with the asymptotic behaviour of a distribution for a function *ψ* of an asymptotically Gaussian random vector. For a sequence of random vectors *X*_*n*_ for which 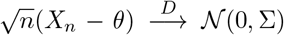, we have that 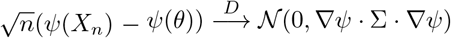. In this context we use the identify

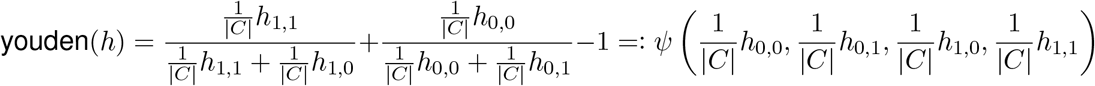

**Fig. 4-d** shows the Youden’s score and its 95% confidence interval for sitka and 6 baseline methods on 3 different real-world datasets. Sitka has a higher score than all competing methods.

### 9.7 Application: assignment of single nucleotide variants

Here we posit an observation probability model for adding single nucleotide variant (SNV) data to an existing phylogenetic tree.

For locus *l* in cell *c*, let 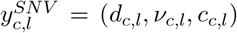 denote the observed SNV data where the total number of reads, the number of reads with a variant allele, and the corresponding copy number are indicated by *d*_*c,l*_, *ν*_*c,l*_, and *c*_*c,l*_ respectively.

We use 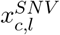 to denote an indicator variable taking the value one if and only if an ancestor of cell *c* harboured a single nucleotide alteration event at locus *l*. This variable is unobserved and the focus of inference in this section. As in the sitka model, we assume a perfect phylogeny structure on these indicator variables, and add an error model to relate 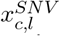 to the observed data while allowing violations of the perfect phylogeny assumption and measurement noise. In the context of single nucleotide data, this is similar to [12]. The parameters of the error model are denoted *θ*^*SNV*^ = (*ϵ*_*FP*_, *ϵ*_*FN*_), where *ϵ*_*FP*_ and *ϵ*_*FN*_ are false positive rate and false negative rates, respectively. Define:

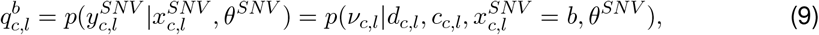

where *d*_*c,l*_ and *c*_*c,l*_ are given inputs. The likelihood probability of cell node *c* is denoted by 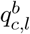, where *b* ∈ {0, 1}. For *b* = 1, 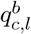 reflects the likelihood of cell *c* being mutated at locus *l*; and for *b* = 0, 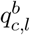 reflects the likelihood of cell *c* not being mutated at locus *l*. For *d*_*c,l*_ = 0, we set 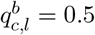.

The probability 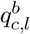 is obtained by marginalizing a mixture of binomial distributions depending on all possible genotype states of locus *l* at cell *c*. Given the copy number *c*_*c,l*_, the possible genotype states are 𝒢= {*A* … *A, AA* … *B, A* … *BB*, …, *B* … *B*}, where each element has a length equal to *c*_*c,l*_. For example, the genotype *AAB* refers to a genotype with one variant allele *B* and two reference alleles *A*. For each genotype state *g*_*i*_, where *i* indexes the elements of **𝒢**, the mean parameter of the corresponding binomial distribution is denoted by 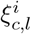 :

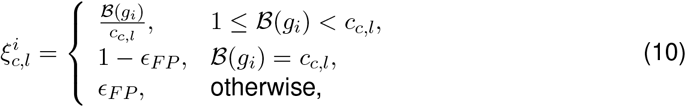

where ℬ(*g*_*i*_) represents the number of variant alleles of genotype *g*_*i*_. Therefore, for *b* = 1,

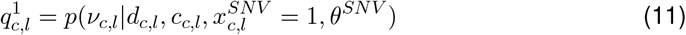

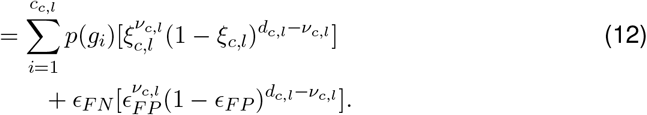

The value of *p*(*g*_*i*_) equals 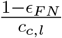, and *ϵ*_*FN*_ represents the error due to mutation loss or tree errors.

If the mutation status of cell *c* at locus *l* is a wildtype (i.e., mutation is not present), then the possible genotype states should not have any variant allele. The only possible genotype state is {*A* … *A*}. The mean parameter of the binomial distribution equals *ϵ*_*FP*_ (false positive rate). Therefore,

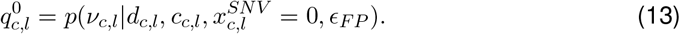

With the proposed probability model for SNVs, we can incorporate both SNV data and CNA data to infer the underlying tree phylogeny in the sitka model. Therefore,

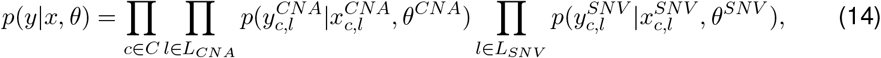

where *C* and *L* are the disjoint set of cells and loci, respectively. In this section, the loci set *L* includes both CNA and SNV traits.

Assume now that we seek to add one locus to an existing tree. We proceed similarly to Section 9.4.3. Equation (4) can be rewritten in the following form:

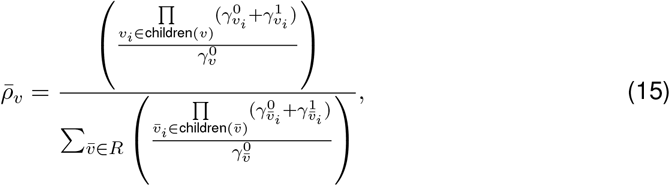

where 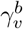, for *b* ∈ {0, 1} is:

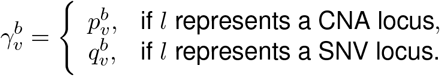

For *v* ∈ *R* = {*v*\*} ∪ *L*\{*l*}, and *b* ∈ {0, 1}, the value of 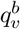 is

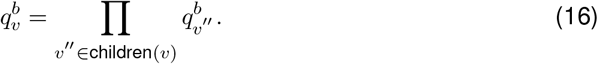

For the cell nodes that are the leaves of the tree 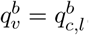.

#### 9.7.1 Detection of SNVs for individual cells

Given a fixed CNA tree (denoted by *t*) and the read counts data (*y*^*SNV*^ denoted by *y* for simplicity), here the goal is to calculate the posterior distribution of 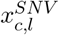, the mutation status of locus *l* at cell *c*, which we denote by *x*_*c,l*_ for simplicity.

The joint probability distribution of *x*_*c,l*_, *y* and *t* can be written as:

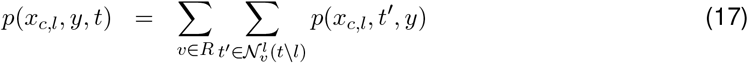

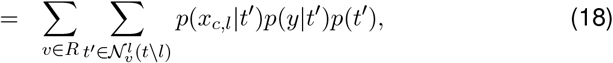

where *R* is the set of all loci nodes in the tree (including the root) excluding locus *l*. The joint probability distribution is calculated as

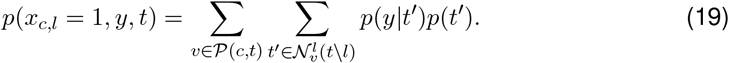

The set 𝒫(*c, t*) denotes all nodes on the shortest path from cell *c* to the root of the tree (including the root and excluding the cell *c* node). An example of the path on an imaginary tree is depicted in **Supplementary Fig. 11**. The nodes coloured in green belong to 𝒫(*c, t*). Therefore, the posterior probability distribution of *x*_*c,l*_ = 1 yields

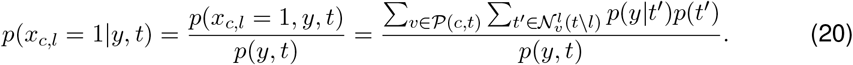

Rewriting Equation (20) assuming uniform probability distribution for *p*(*t*′) yields:

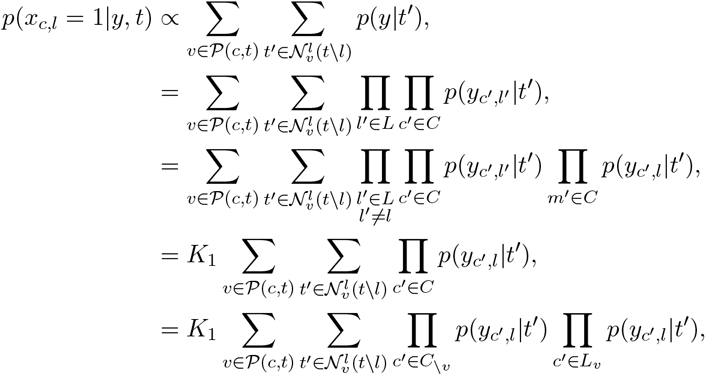

where *N* denotes the set of all trait nodes, *C* denotes the set of all cell nodes, *C*_*v*_ denotes the cells that are a descendant of node *v*, and *C*_\*v*_ denotes the cells that are a not descendant of node *v*. The product of the likelihood contributions for non-descendant nodes can be calculated by taking the product of 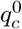for all cells, divided by the ones that are descendant of *v*:

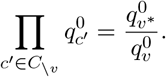

Therefore:

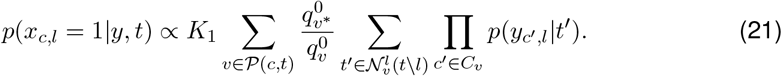

The likelihood contribution of descendant cells can be re-indexed by a binary vector **b** = (*b*_1_, *b*_2_, …, *b*_*k*_), where *b*_*i*_ 0, 1, and *b*_*i*_ = 1 if the child *v* is to be moved into a child of the node *l*. The value of *k* denotes the number of children of *v*. The *i*\*th child of *v* which is on the path from node *v* to cell *c* is called 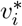. This implies *b*_*i*\*_ = 1 (See **Supplementary Fig. 11**). Therefore:

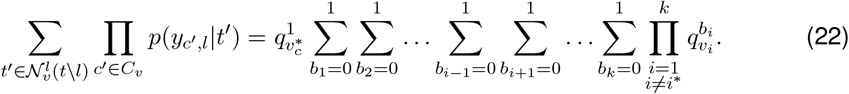

Rewriting Equation (21) using Equation (22) yields:

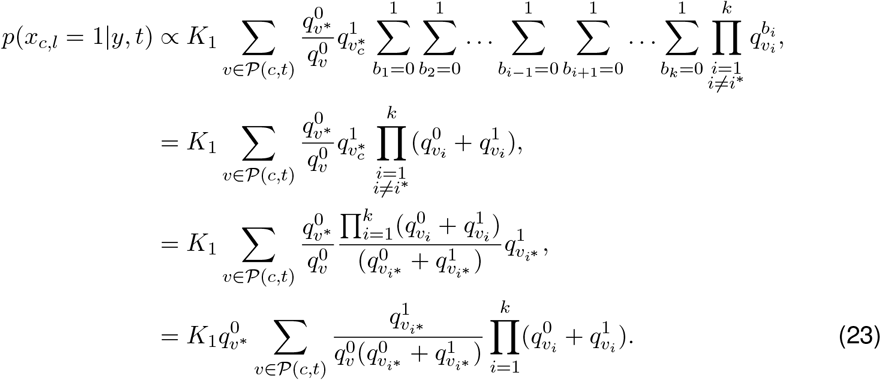

### 9.8 Computational complexity of the SNV calling algorithm

The computational complexity of Equation (23) is *O*(|*C*| · |*L*|) with |*C*| the number of cells and |*L*| the number of loci. In order to reduce the complexity of calculating *p*(*x*_*c,l*_ = 1 |*y, t*) for each locus and cell, 𝒫′(*c, t*) is defined to denote the nodes sitting on the path from root to cell *c*, excluding the root node and including the cell *c* node. Then,

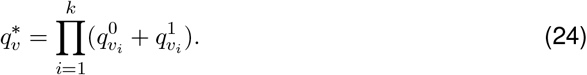

Therefore,

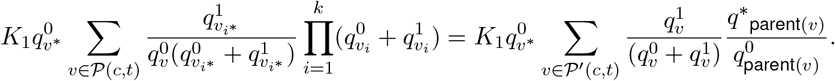

Calculating *p*(*x*_*c,l*_ = 1|*y, t*) with a recursive approach reduces the complexity from *O*(|*C*||*L*|) to *O*(|*C*| + |*L*|), where as in the last section *L* is the union of SNV and CNA loci.

## List of Supplementary Figures

**Supplemental Figure 1.**
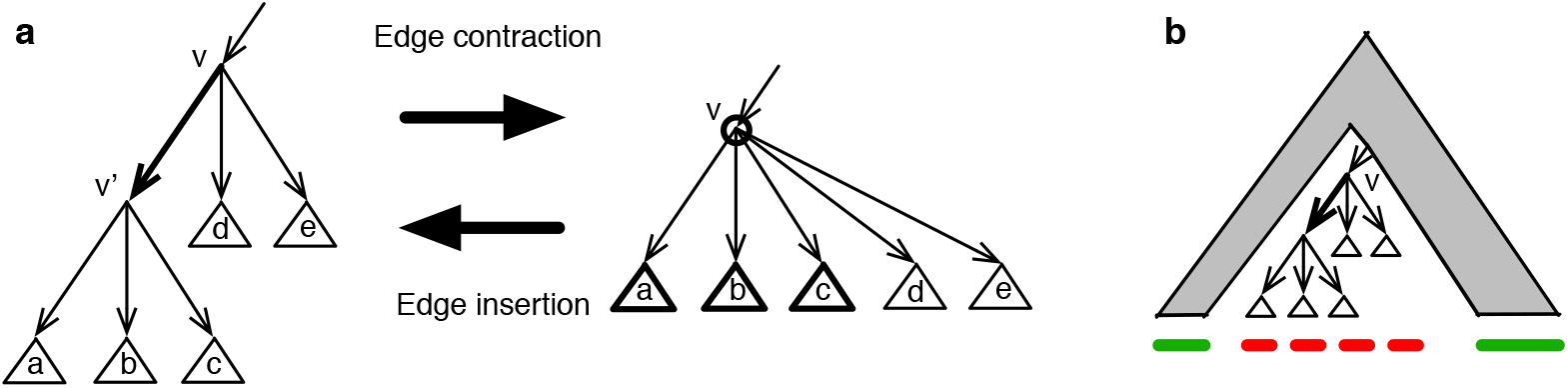
(**a**) Reading from left to right: the interpretation of removing a column in the matrix *x* is to perform contraction of an edge corresponding to a locus shown in bold. Reading from right to left: the interpretation of inserting back a column while assigning new binary values is an edge insertion. The circled node *v* refers to Step 1. The subtrees in bold refer to those selected in Step 2. The edge in bold, the one introduced in Step 3. (**b**) Decomposition used for the recursion of Section 9.4.3.

**Supplemental Figure 2.**
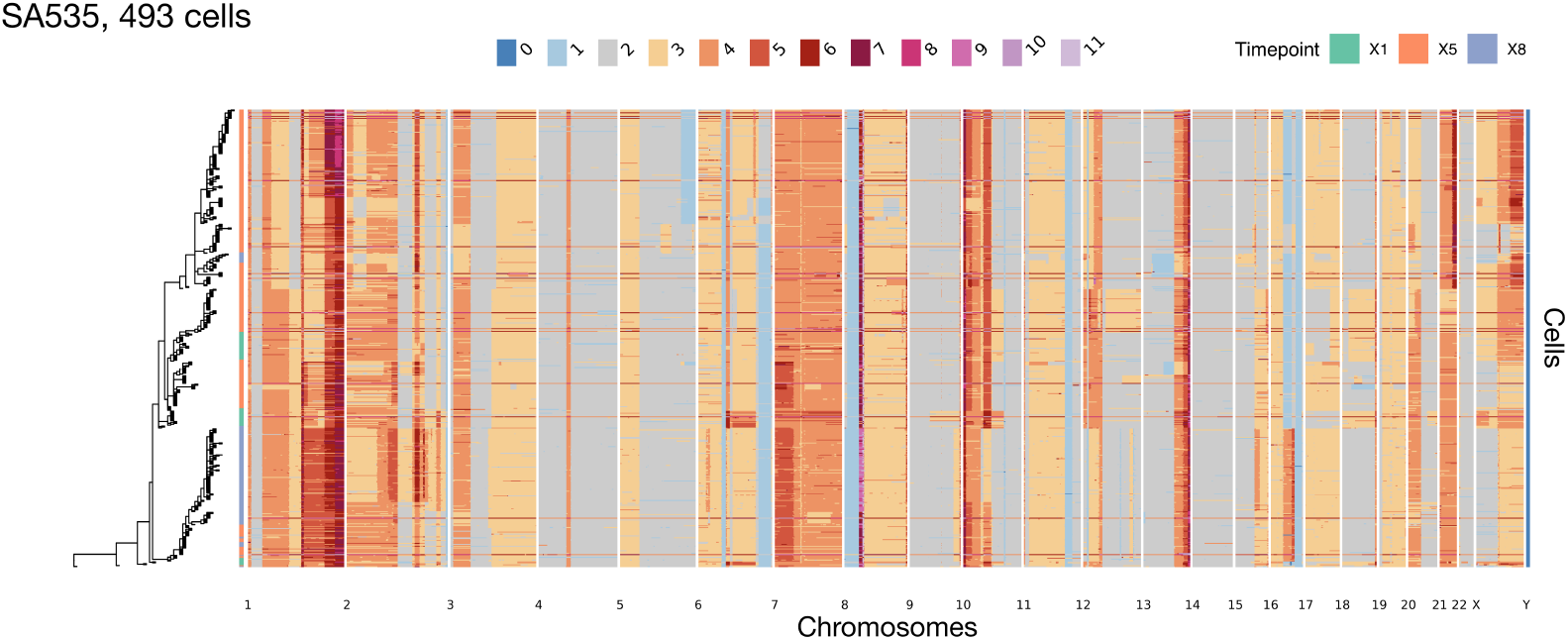
Phylogenetic tree and CNA profile heatmap for the SA535 dataset. The rows of the heatmap are sorted according to the placement of cells on the phylogenetic tree. The columns of the heatmap are sorted by their genomic position.

**Supplemental Figure 3.**
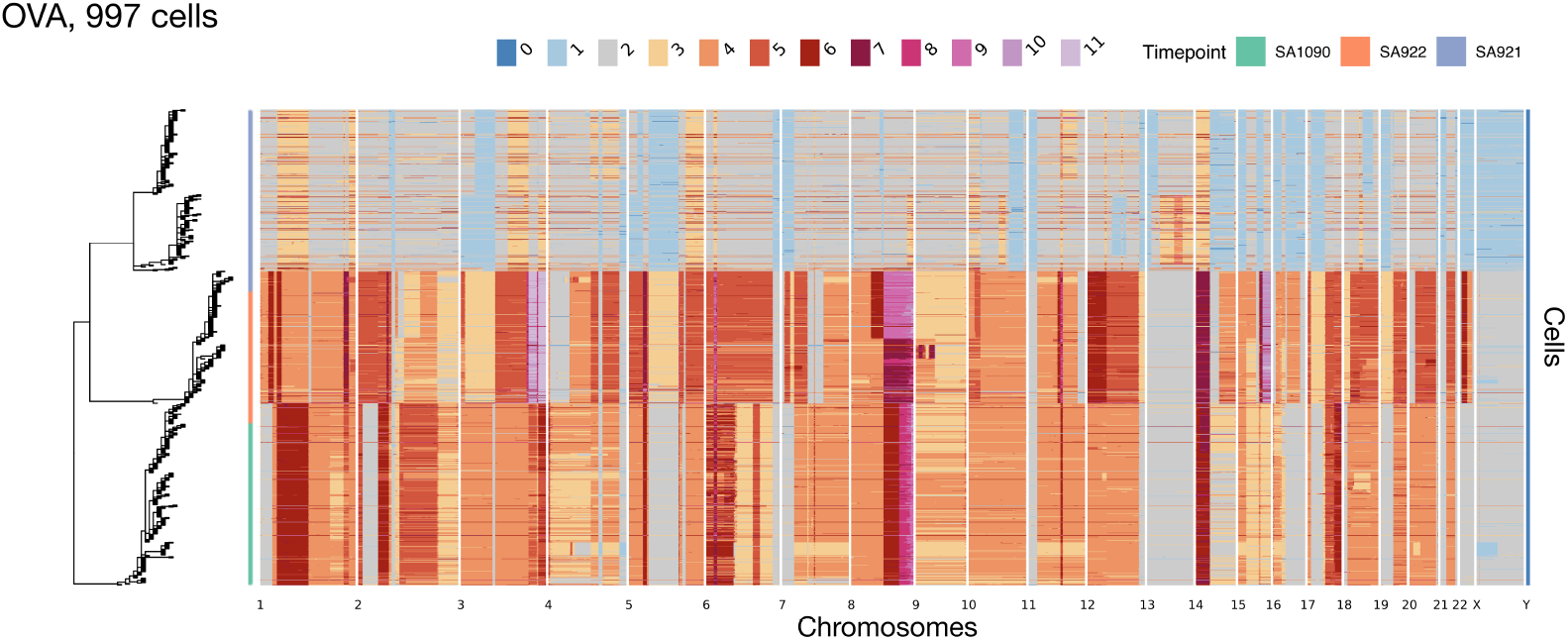
Phylogenetic tree and CNA profile heatmap for the *OV A* dataset. The nearly diploid cells with the loss of heterozygosity on chromosome X are from SA1090. The cells with an amplification on chromosome 22 are from SA922. The rest belong to SA921.

**Supplemental Figure 4.**
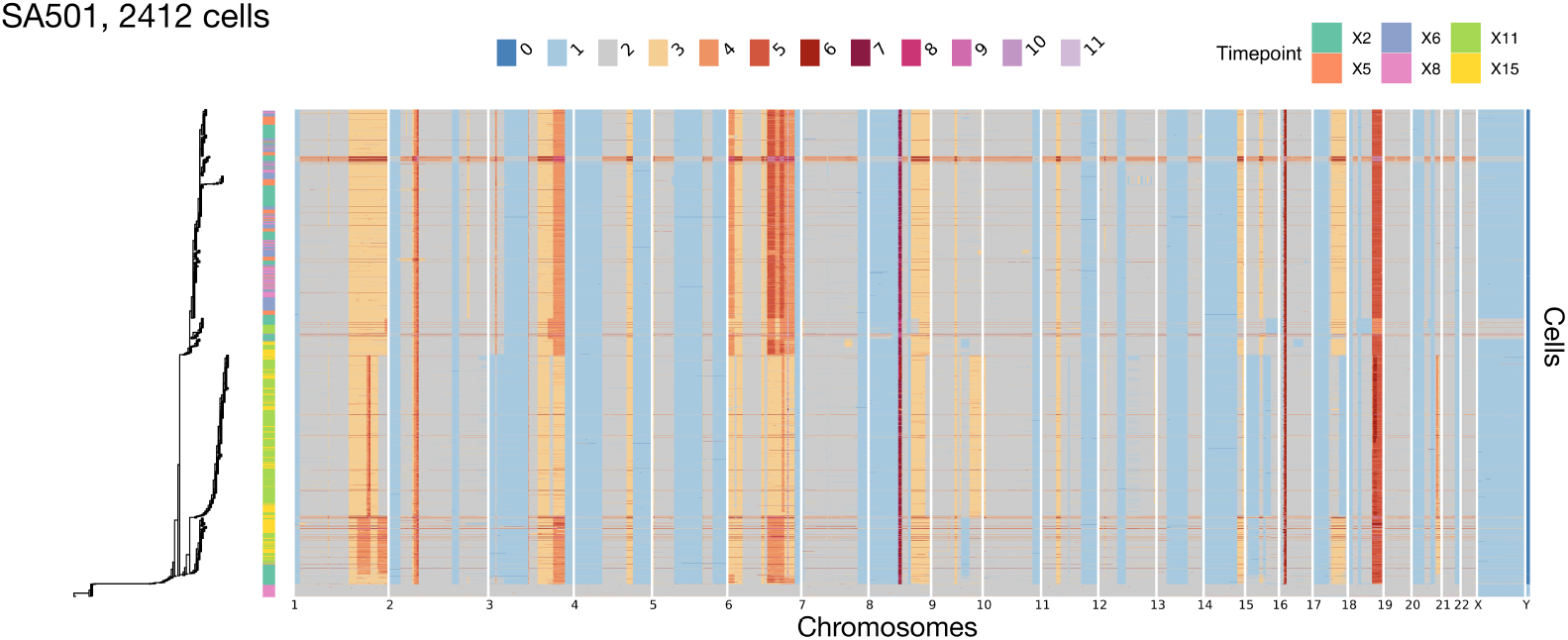
Phylogenetic tree and CNA profile heatmap for the SA501 dataset. Note that the diploid cells at the bottom of the heatmap are control cells that were included in the experiment.

**Supplemental Figure 5.**
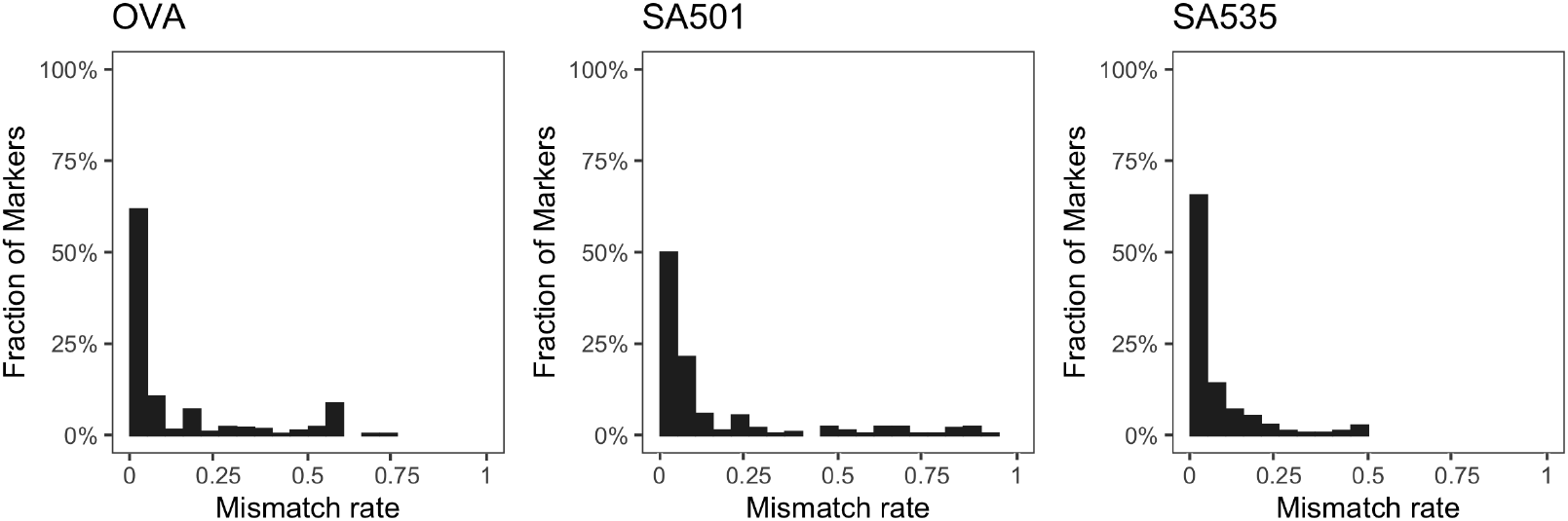
the distribution of mismatch rate defined as the fraction of cells that have a mismatch between the inferred and jitter-fixed value of a marker.

**Supplemental Figure 6.**
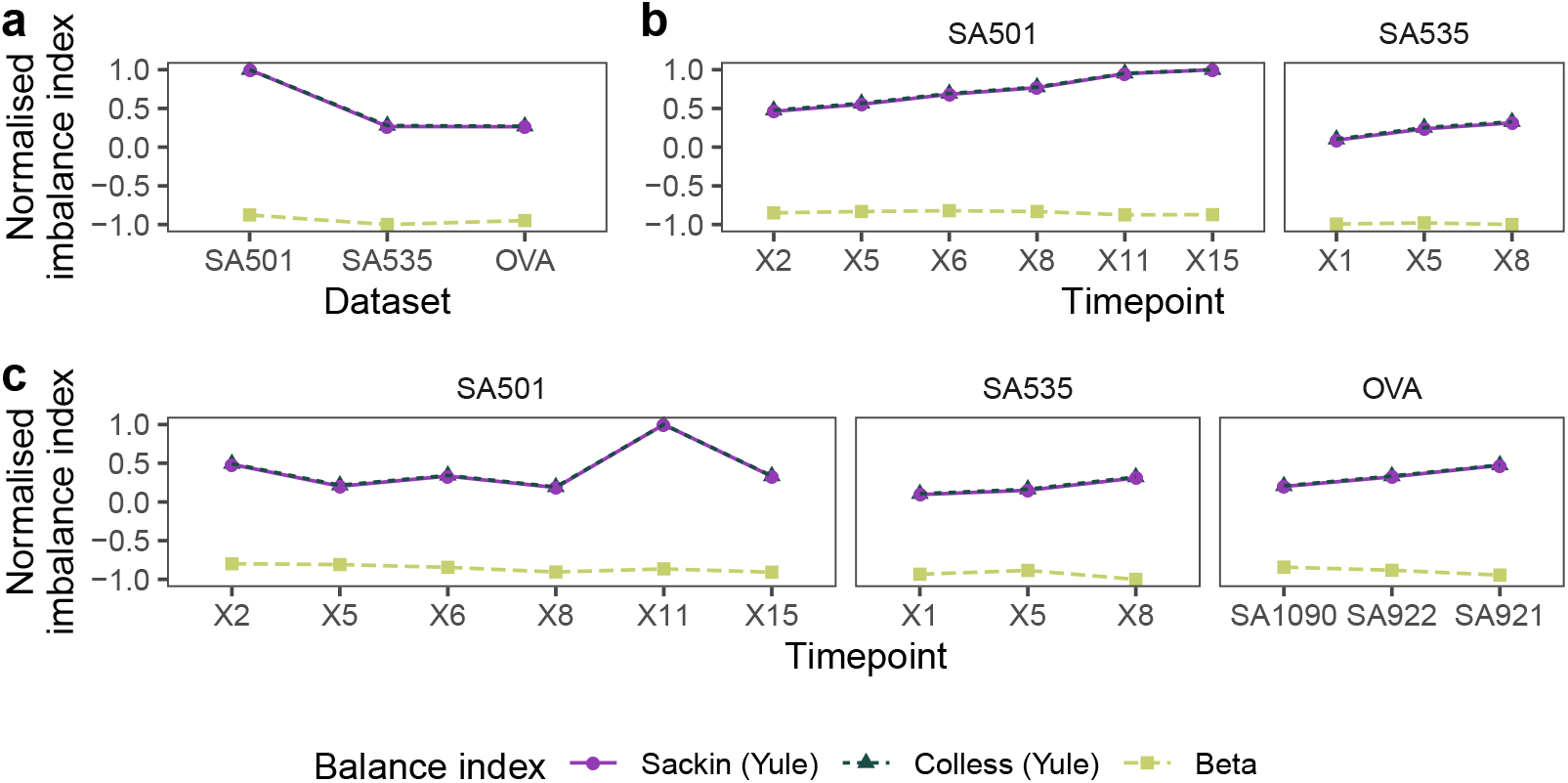
**(a)** Tree imbalance index where zero indicates that the tree is consistent with one simulated from a Yule model (completely balanced) and positive values indicate deviation from the Yule model (more imbalanced). For ease of plotting, each balance index is normalised by the absolute value of the maximum estimated statistic among all samples. Cumulatively adding more timepoints (**b**), or for the maximal subtree comprising cells of a specific timepoint (**c**).

**Supplemental Figure 7.**
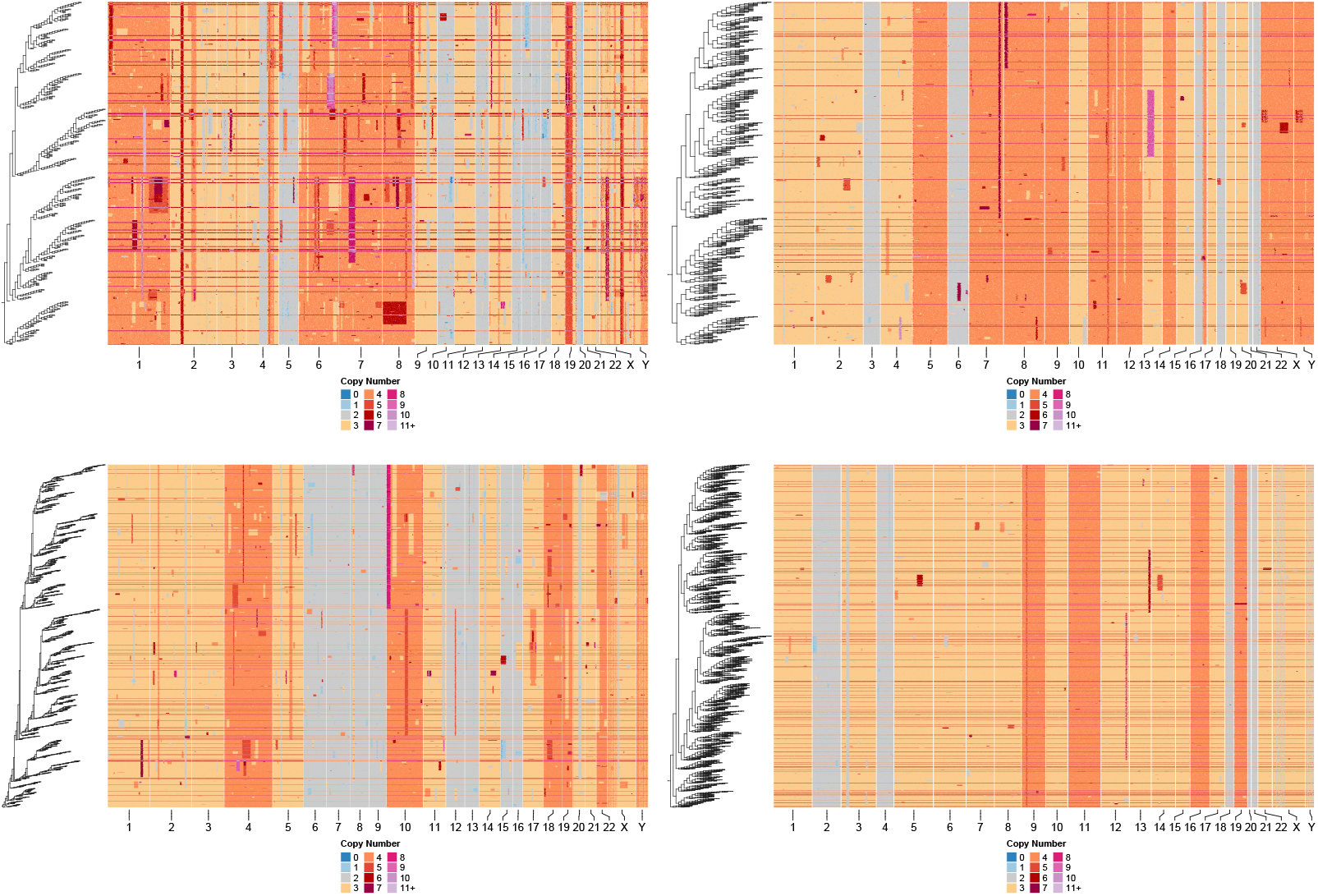
Synthetic datasets simulated from the GRFBS model. The number of cells, number of loci, and *β* values are are (500 × 800, − 0.83), (1000 × 400, − 0.1), (2000 × 800, − 0.9), (3000 × 400, − 0.48) for the upper left, upper right, lower left, and lower right figures respectively.

**Supplemental Figure 8.**
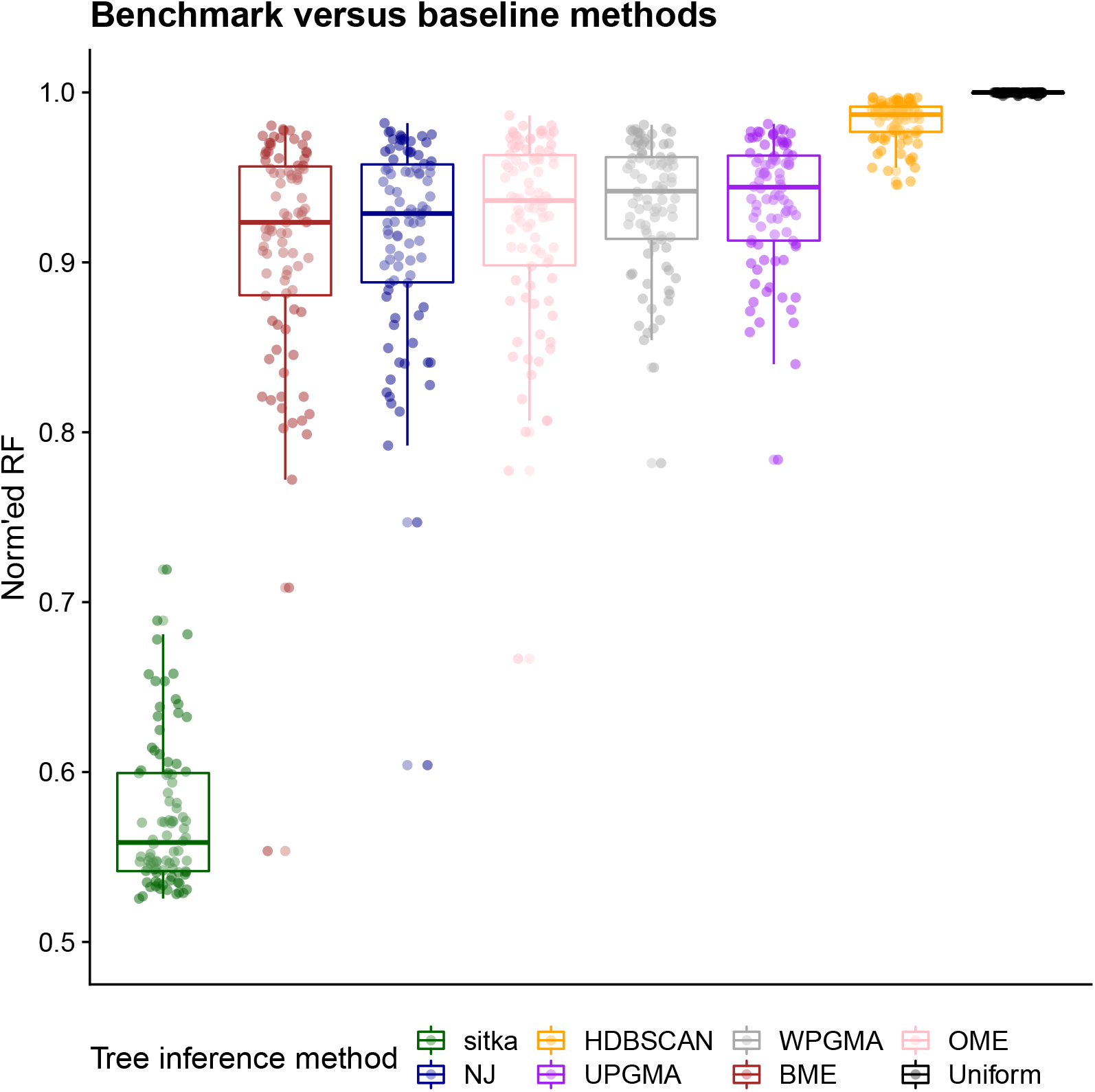
Tree reconstruction evaluation using a normalized Robin-son–Foulds metric on synthetic datasets from *S*90, simulated from Beta-splitting processes. Here normalization is done by dividing the RF distance of each inference method by the worst performer per dataset.

**Supplemental Figure 9.**
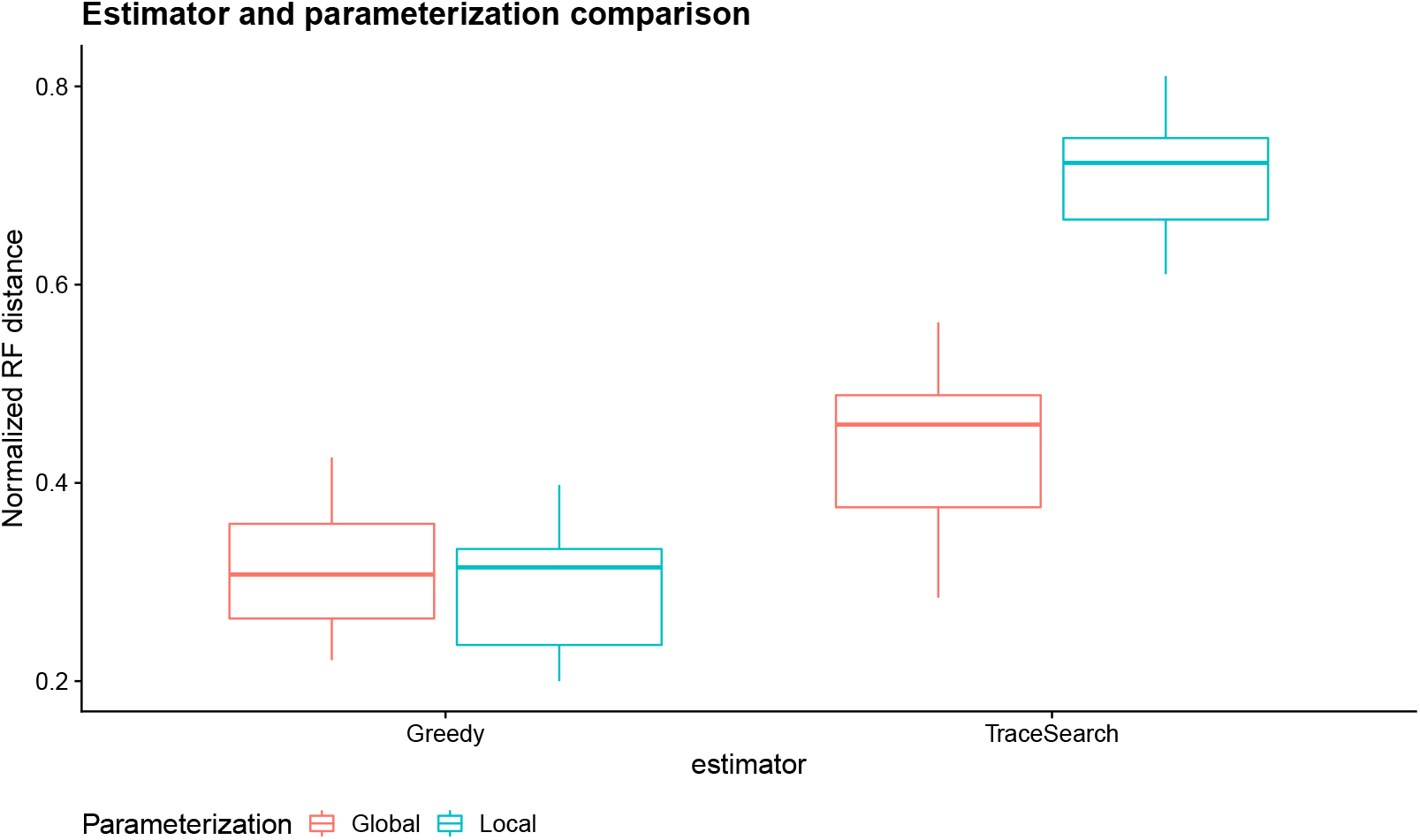
A model and estimator comparison based on tree reconstruction accuracy for datasets from *S*10. For each dataset, inference was performed on both the globally- and locally-parameterized model. Both the greedy and trace search estimates were computed for each inference result.

**Supplemental Figure 10.**
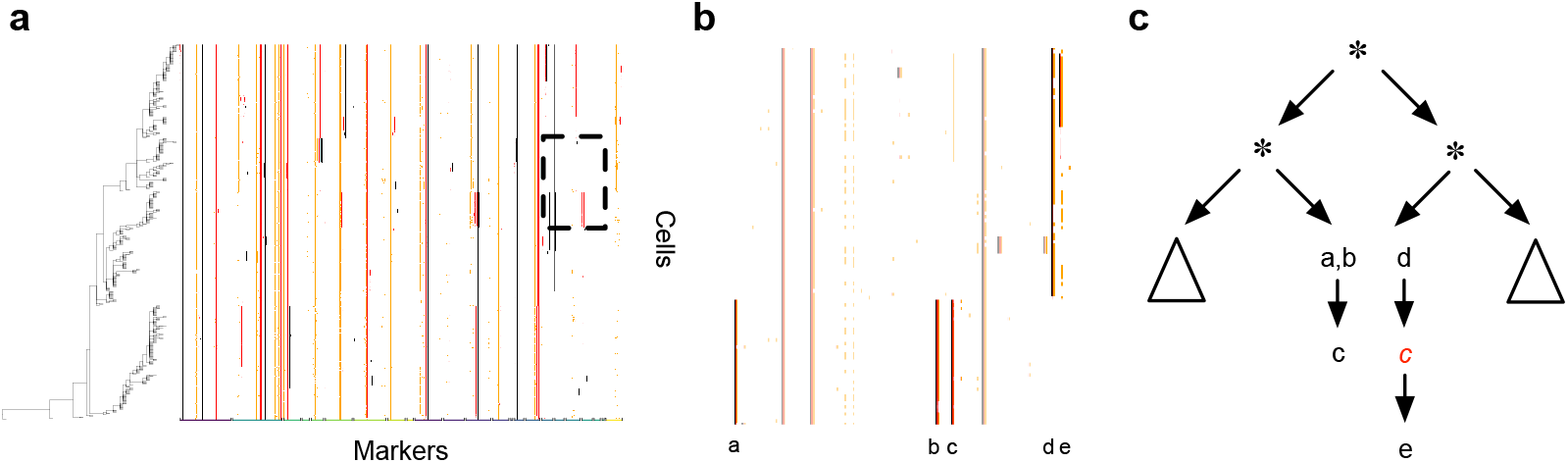
A locus in *SA*535 that violates the perfect phylogeny assumption. The method is robust to this violation and correctly allocates the other relations (positive *x* entries shown in black in the matrix; positive *y* entries shown in orange; posterior marginals *m* in shades of red). **(a)** the part of the tree where the violation occurs, magnified in **(b)**, where the short band of orange not matched with a black band is indicative of the violation. **(c)** the corresponding tree showing the two points of the tree where the two events collocated in the same bin occurred in the tree. The sister clades of a,b and d suggest that this is a case of two insertion events rather than a loss.

**Supplemental Figure 11.**
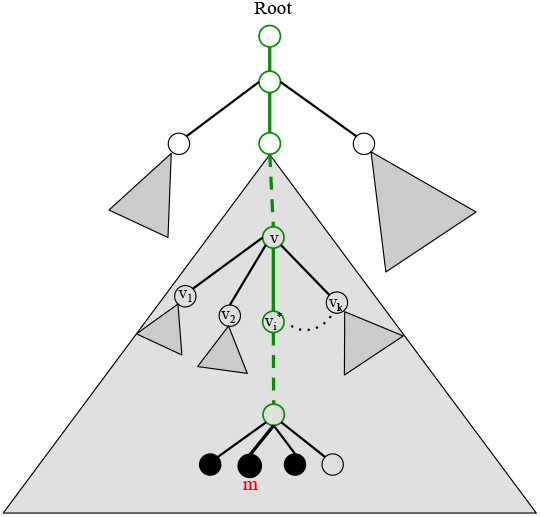
A schematic view of the underlying tree inferred from CNA and SNV loci across multiple cells. Black and white nodes represent cells and loci, respectively. The grey triangle represents a subtree rooted at a node. It includes all of the nodes and edges in the subtree.

**Supplemental Figure 12.**
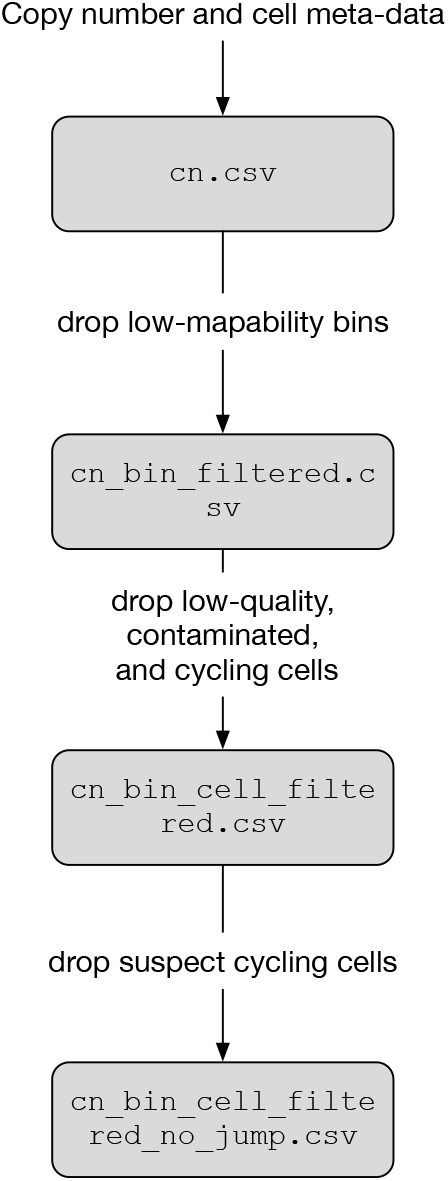
Filtering the CNA data for tree inference.

**Supplemental Figure 13.**
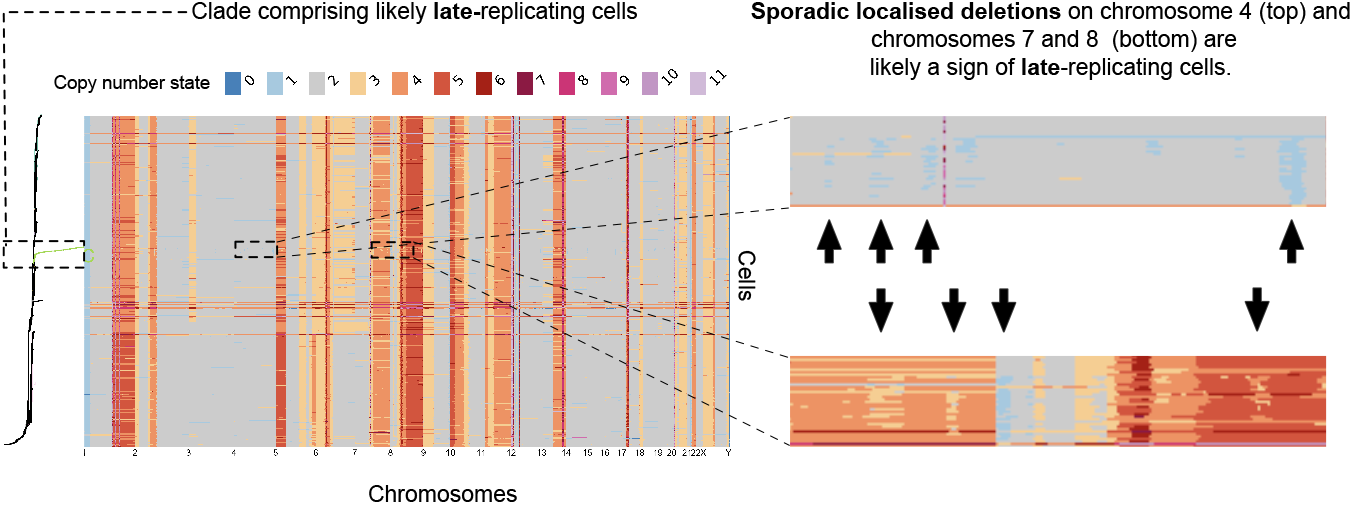
An example of replicating cells. Note the scattered localised deletions. This heatmap is from a HER2+ PDX line. These late replicating cells form a *finger* like clade in the tree. The top inset shows chromosome 4 while the bottom inset spans chromosomes 7 and 8.

**Supplemental Figure 14.**
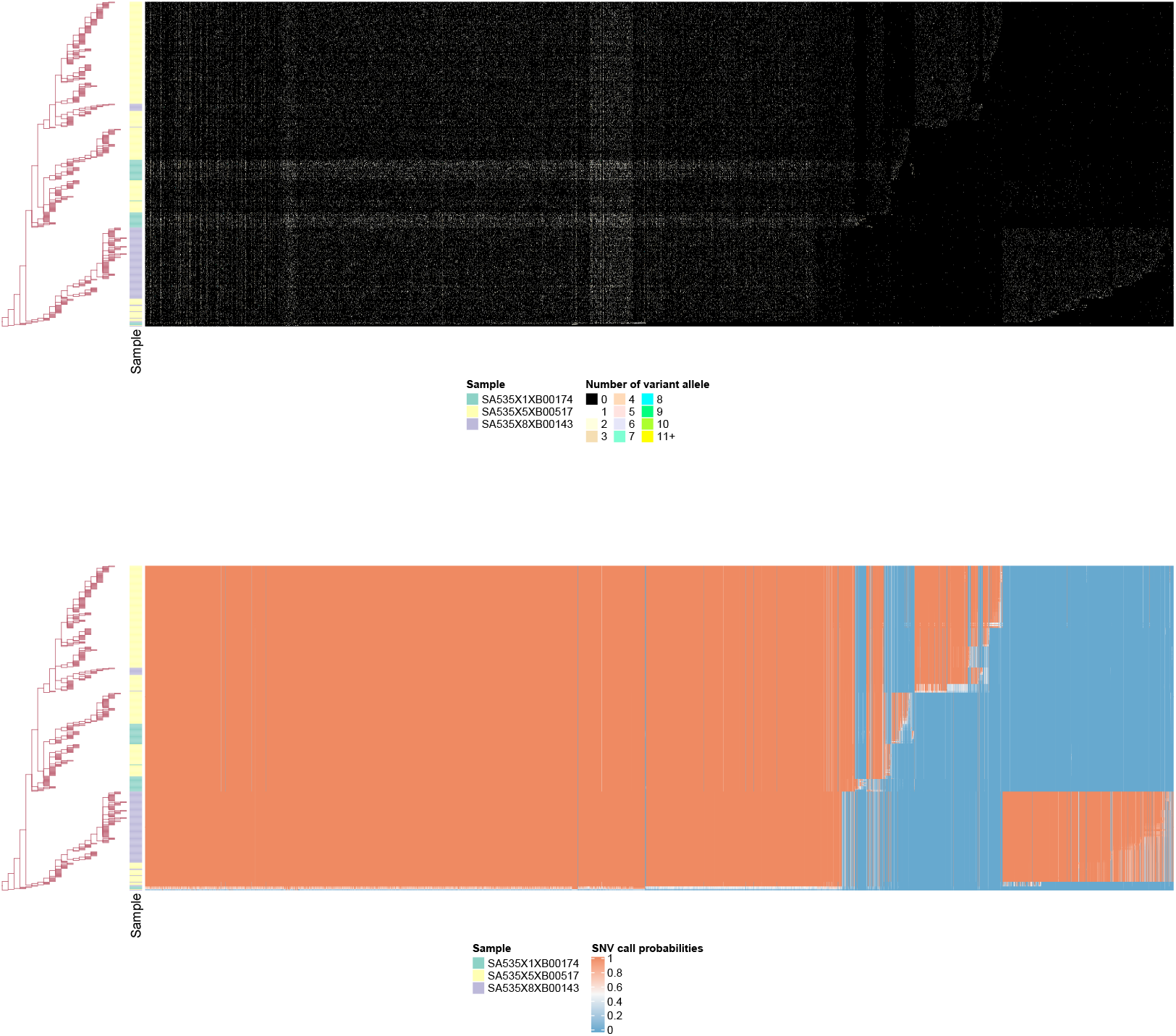
SNV variant reads data and SNV call probabilities for SA535 dataset beside the underlying phylogenetic tree.

**Supplemental Figure 15.**
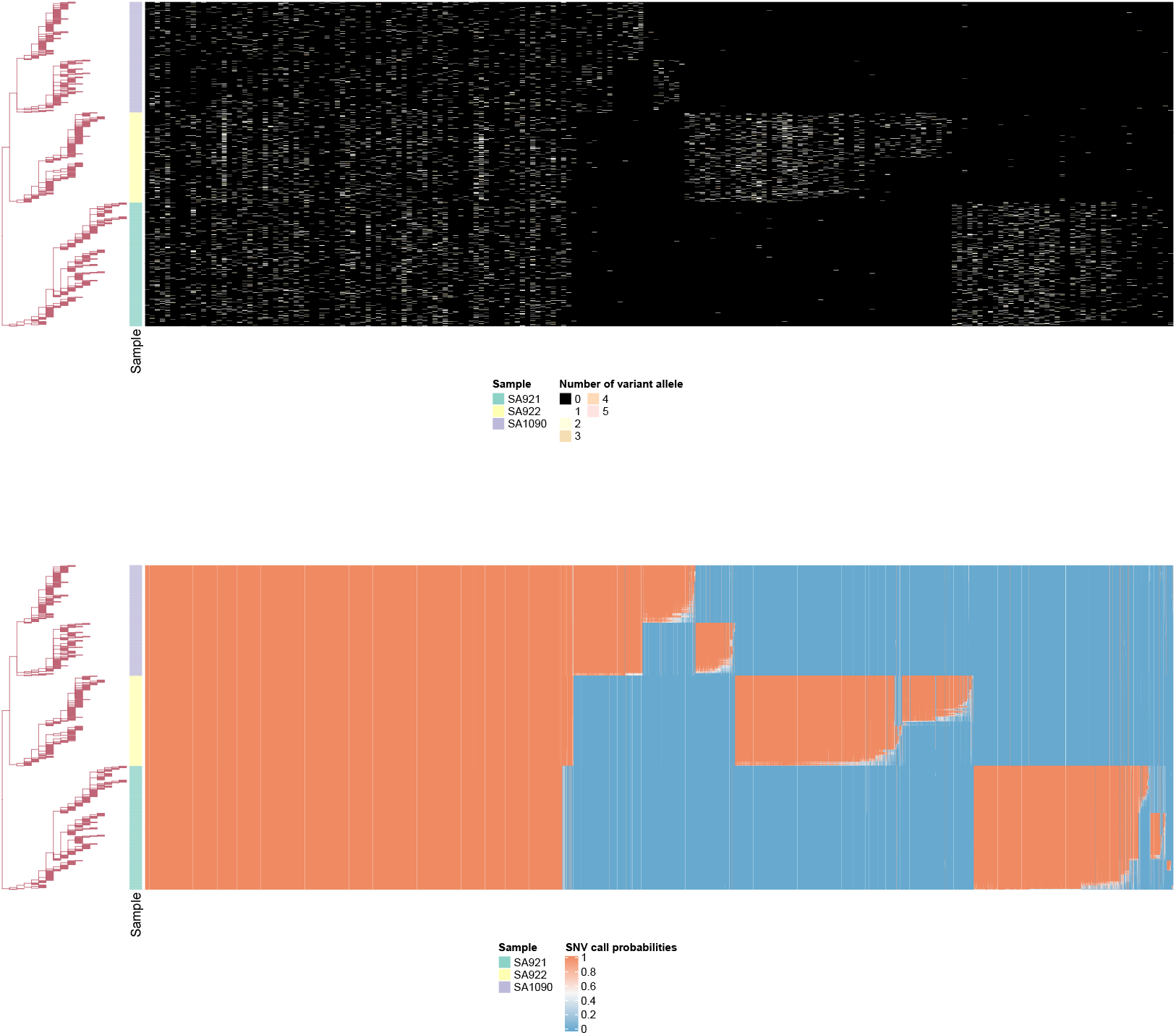
SNV variant reads data and SNV call probabilities for OVA dataset beside the underlying phylogenetic tree..

**Supplemental Figure 16.**
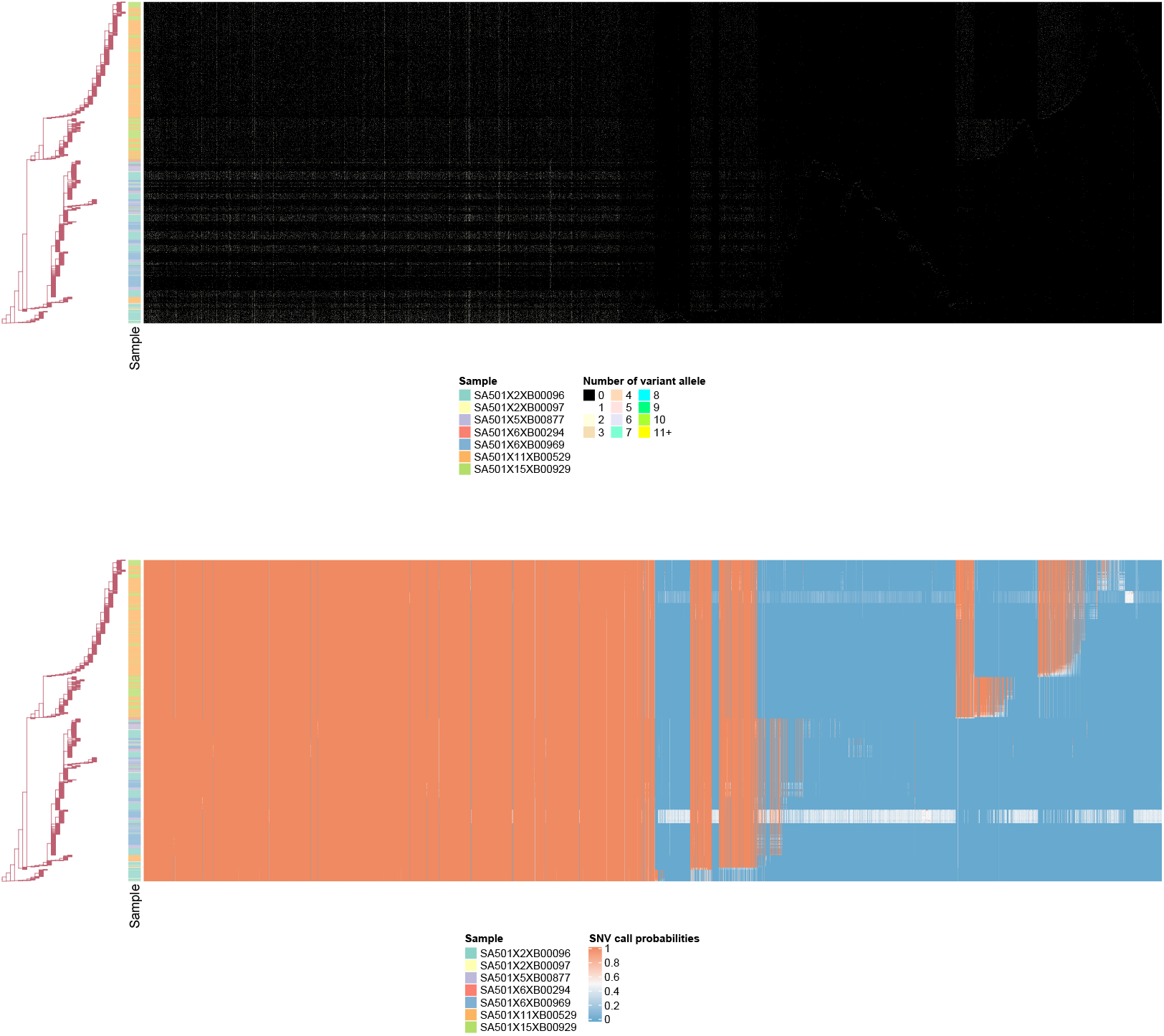
SNV variant reads data and SNV call probabilities for SA501 dataset beside the underlying phylogenetic tree.

**Supplemental Table 1.** Summary of real-world datasets used. `final` is the final number of cells after all filters except for `!lmr` are applied. `final` additionally filters out `lmr` cells, those that have total mapped reads fewer than 500,000. Abbreviations used are tp: time point; qual. : quality; !sphase: not sphase; !lmr: not low mapped reads.

| Dataset | parameter | value |
| --- | --- | --- |
| Real datasets | engine | PT |
| Real datasets | globalParameterization | true |
| Real datasets | fprBound | 0.1 |
| Real datasets | fnrBound | 0.5 |
| Real datasets | nChains | 1 |
| Real datasets | nScans | 1000 |
| Real datasets | nPassesPerScan | 1 |
| Real datasets | thinning | 1 |
| Real datasets | burnin fraction | 0.5 |
| *S90* | engine | PT |
| *S90* | globalParameterization | true |
| *S90* | fprBound | 0.5 |
| *S90* | fnrBound | 0.5 |
| *S90* | nChains | 1 |
| *S90* | nScans | 20000 |
| *S90* | nPassesPerScan | 1 |
| *S90* | thinning | 1 |
| *S90* | burnin fraction | 0.5 |
| *S10* | globalParameterization | true, false |
| *S130* | globalParameterization | true |
| *S10,S130* | engine | PT |
| *S10,S130* | fprBound | 0.1 |
| *S10,S130* | fnrBound | 0.5 |
| *S10,S130* | nChains | 8 |
| *S10,S130* | nScans | 5000 |
| *S10,S130* | nPassesPerScan | 10 |
| *S10,S130* | thinning | 1 |
| *S10,S130* | burnin fraction | 0.5 |

**Supplemental Table 2**. Inference settings used for each dataset.

## Footnotes

1 Authors of [42] argue the GBFBS model is capable of realizing topologies comparable to that of the original Beta-splitting model. When *β* = *α* → −1, trees are totally imbalanced; when *β* = *α* →*∞*, trees are perfectly balanced.

2 We used the R packages [47, 48] for simulation.

