## Supplemental Text for "Cancer phylogenetic tree inference at scale from 1000s of single cell genomes"

#### 1 Pre-processing

##### 1.1 HMMcopy pre-processing

Corrected CNA states from `HMMCopy` are stored in `cn.csv.gz` and this file is the input to our preprocessing pipeline. For this work the data was stored in the cloud (Microsoft Azure) and the `scgenome` API was used to access and download the data. Please see `https://github.com/shahcompbio/scgenome` for documentation. The `scgenome` API ensures that only cells with the correct `sample_ids` are selected, removes control cells and cells that have fewer than 10,000 mapped reads.

##### 1.2 Filtering low-mappability bins

Some copy number bins are located at parts of the genome where sequencing is difficult, for example due to inaccessibility of the genome at that position. This is reflected in their mappability score. We filter the CNA matrix to keep high-map-ability bins `cn_bin_filtered.csv.gz`. In this work we use a cutoff threshold of `map >=.99` that yields $4375/6206$ or $0.705\%$ of the bins. The list of kept bins is identical across all datasets.

*Equal contribution

#### 1.3 Filtering low-quality cells

In this step, a second round of quality control is done. Cells with a *quality score*, as defined in [**laks_directlibrary_2019**], of over 0.75 or higher are kept, while the other ones which are suspected to be contaminated (e.g., mouse cells) or cycling cells are removed. This results in `cn_bin_cell_filtered.csv.gz`.

#### 1.4 Filtering cells with excess CNA changes

Some cells show a "jumpy" CNA profile in which there are too many copy number changes. It appears that these cells are either in early or late stages of division and were missed by the S-phase classifier. The CNA profile of early replicating cells is patterned by seemingly scattered focal amplifications while the late replicating cells show scattered focal deletions. Note that not all parts of the genome duplicate at the same time upon cell division. Regions that start duplicating later will show as having focal deletions in cells captured at their later replicating stage; these regions would have not started to duplicate by the time the sample was prepared for sequencing. These scattered patterns 13 do not directly reflect the evolutionary history of the cells and are detrimental to phylogenetic tree inference.

Here we rank cells by the number of changes in their copy number states (a change is measured between consecutive bins) and pick the $90\%$ percentile. The file `cn_bin_cell_filtered_no_jump.csv.gz` contains the integer copy number state with the final list of cells and genomic bins. An example input matrix is shown in **Supplementary Fig. 3-a** where the integer copy numbers are coloured coded in a heatmap. Attrition rate due to filtering of cells is shown in Table 1.

### 2 Baseline methods

Here we give a brief description of the baseline methods to which we compare sitka. `UPGMA` [1], `WPGMA` [1] and Neighbour Joining (NJ) [2] are all distance-based phylogenetic inference methods, that is, they use the input data to first compute a similarity matrix between single-cells and then proceed to construct an agglomerative clustering in an iterative process. HDBSCAN [3] on the other hand is a heuristic that, roughly speaking, computes the minimum spanning tree from a low dimensional representation of the CN matrix. MEDALT [4] is another distance based method that uses the Chu–Liu's algorithm to compute a directed minimum spanning tree from the matrix of minimum edit distance. Medicc2 [5] uses a finite state transducer to model copy number evolution over time, taking into account allele specific and whole genome duplication events. MrBayes [6] is a Bayesian phylogenetics framework that implements multiple evolutionary models and uses MCMC to approximate the posterior distribution of trees and model parameters.

### 3 Tree shape statistics

All three trees from the three real data experiments are imbalanced [7] relative to a Yule model (**Supplementary Fig. 6-a**). Unbalanced tree topologies appear and are expected in adapting populations [8]. In $SA535$ and $OV2295$ the sample subtrees become more balanced over time and post-relapse respectively. In contrast, $SA501$ exhibits a decrease in balancedness, except timepoint X11, where a marked increase in imbalance is observed (**Fig. 6**-**b,c**).

sitka inferred trees are not dichotomous therefore we first resolve multichotomies into dichotomies. As there are multiple ways to resolve multichotomies, we do this 100 times, resolving multichotomies uniformly at random. We then compute the balance statistics on each resulting dichotomous tree and report the average in **Fig. 6**-**a-c**. We report three balance statistics, namely Sackin, Colless, and Beta [9].

The Sackin [10] and Colless [10] statistics are both measures of imbalance of a rooted phylogenetic tree. The former is the sum of the depth of the leaves, while the latter is the sum of the absolute values of the difference between the number of descendent leaves of the left and the right child of each internal node. The Yule [10] model is a probabilistic model for bifurcating phylogenetic trees. Under this stochastic model, a phylogenetic tree with N leaves is generated by an iterative process: given rooted tree with 2 leaves, pick a leaf node uniformly randomly, and attach two new leaf nodes to it until the tree has N nodes.

We measure the change in the balance of the tree over time in two ways: (i) starting with the first timepoint and progressively adding more timepoints **Fig. 6**-**b**), (ii) in individual timepoints **Fig. 6**-**c**). In the former, we start with the maximal subtree $\tau_{\text{max}}(X_{t_0})$ for cells in the first timepoint ($X_{t_0}$), then compute the maximal subtree that contains the first two timepoints $\tau_{\text{max}}((X_{t_0}, X_{t_1}))$, and continue until all timepoints are included. We report the imbalance statistic for each subtree constructed in this process. In the latter, for each timepoint $X_t$, we find the maximal subtree $\tau_{\text{max}}(X_t)$ that contains all cells from timepoint $X_t$, and report the imbalance index for it.
